# Enhanced Feature Selection for Microbiome Data using FLORAL: Scalable Log-ratio Lasso Regression

**DOI:** 10.1101/2023.05.02.538599

**Authors:** Teng Fei, Tyler Funnell, Nicholas R. Waters, Sandeep S. Raj, Keimya Sadeghi, Anqi Dai, Oriana Miltiadous, Roni Shouval, Meng Lv, Jonathan U. Peled, Doris M. Ponce, Miguel-Angel Perales, Mithat Gönen, Marcel R. M. van den Brink

## Abstract

Identifying predictive biomarkers of patient outcomes from high-throughput microbiome data is of high interest, while existing computational methods do not satisfactorily account for complex survival endpoints, longitudinal samples, and taxa-specific sequencing biases. We present `FLORAL` (https://vdblab.github.io/FLORAL/), an open-source computational tool to perform scalable log-ratio lasso regression and microbial feature selection for continuous, binary, time-to-event, and competing risk outcomes, with compatibility of longitudinal microbiome data as time-dependent covariates. The proposed method adapts the augmented Lagrangian algorithm for a zero-sum constraint optimization problem while enabling a two-stage screening process for extended false-positive control. In extensive simulation and real-data analyses, `FLORAL` achieved consistently better false-positive control compared to other lasso-based approaches, and better sensitivity over popular differential abundance testing methods for datasets with smaller sample size. In a survival analysis in allogeneic hematopoietic-cell transplant, we further demonstrated considerable improvement by `FLORAL` in microbial feature selection by utilizing longitudinal microbiome data over only using baseline microbiome data.

## 1 Introduction

Advances in computational approaches for analyzing metagenomic data have substantially improved our understanding of the relationships between the human microbiota and environmental exposures, health conditions, treatment responses, and patient survival. Discovery of microbiome compositions predictive of human disease or treatment outcomes provide opportunities for therapeutic intervention[1]. At the same time, the rapidly evolving technology and quickly accumulating amount of available microbiome data over the past decade have motivated computational biologists and biostatisticians to develop robust analytical approaches to detect associations between microbiota and factors of interest, while avoiding false-positives [2, 3].

Allogeneic hematopoietic cell transplants (allo-HCT) provides a paradigm for understanding the importance of microbiome composition in clinical outcomes. While provided to patients with curative intent, high-dose chemotherapy prior to the transplant causes severe damage to gut microbiota, which further increases the risk of life-threatening gut inflammation, opportunistic infections, and malnutrition. Therefore, it is of high interest to monitor and study the association between the microbial profiles and the corresponding patient outcomes, which are commonly coded as continuous, binary, time-to-event, or competing risks outcomes [4].

To fulfill the need of identifying microbial biomarkers, differential taxa abundance analysis approaches have been applied to compositional microbiome data [5]. As the sequencing depth may heavily vary across samples, one should account for both observed count and sequencing depth (i.e. total read count per sample) to facilitate a standardized quantification of a taxon of interest across samples. One naive approach is to perform the two-sample Wilcoxon rank-sum test for the observed relative taxon abundance (count divided by sequencing depth). More sophisticated strategies include applying multi-stage Wilcoxon tests and linear discriminant analysis (`LEfSe` [6]), modeling high presence of zero counts (`metagenomeSeq` [7], `ANCOM-II` [8], `ANCOM-BC` [9]), direct modeling of count data (`ALDEx2` [10], `corncob` [11]), and performing permutation tests (`LDM` [3, 12, 13]).

While the above differential abundance (DA) testing methods are useful, there are some important limitations. Typically, the DA methods perform multiple hypothesis testing followed by p-value adjustment, where taxon-outcome associations are assessed in a univariable manner without accounting for other taxa, which tends to inflate the number of selected taxa. In addition, taxa selection is determined by a chosen p-value threshold, where the choices of 0.2, 0.1 and 0.05 have been widely reported without consensus [14–16], potentially contributing to reproducibility issues in microbiome research [**citations**]. Moreover, the majority of DA methods lack utilities of handling time-to-event response variables and longitudinal microbiome data, compromising the best use of data by performing comparisons across binary “event” and “non-event” groups without accounting for follow-up [17]. Furthermore, taxon-specific sequencing bias may disrupt the rank of relative abundances across samples [18], suggesting methods based on relative abundance or sequencing depths may suffer potential performance loss.

As an alternative approach to identifying the taxa-outcome association, penalized log-ratio regression (or log-ratio lasso) models were derived from classic compositional data regression [19], treating ratios between microbial features as predictors, with linear [20–23], binary [21–23], or time-to-event [21] outcome variables. Since there are 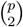 unique pairwise ratios out of *p* taxa, computationally efficient algorithms with zero-sum constrained loss functions [20–22] were widely established, avoiding direct enumeration of ratios [23]. In addition, a two-step variable selection scheme was proposed to further reduce the false discovery rate [22]. Unlike the DA methods, log-ratio lasso regression assesses taxa-outcome associations in multivariable models, conducts variable selection using more objective criteria based on cross validations, naturally incorporates various types of response variables including time-to-event, and effectively circumvents the potential issues introduced by taxa-specific sequencing biases [18]. Nevertheless, currently available software packages (`zeroSum` [21], `logratiolasso` [22]) have not comprehensively implemented all previously developed features for various outcome types or variable selection strategies. Moreover, the existing methods were not developed to incorporate complex outcomes such as competing risks [4, 24], or time-dependent microbial predictors, which have already been widely available in large-scale longitudinal clinical studies.

Here we propose `FLORAL` to perform linear, logistic, Cox proportional hazards [25], and Fine-Gray proportional subdistributional hazards [26] log-ratio lasso regression and subsequent feature selection for high-dimensional compositional data (**Fig.1**). We develop a unified loss function framework that can easily adapt various types of outcome variables (**Fig.1A**). Instead of enumerating 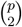 possible pairs of taxa, the proposed algorithm works on the *p*-dimensional covariate space as facilitated by the zero-sum constraint, which only requires affordable computing memory. To accommodate longitudinal microbiome data, `FLORAL` enables time-dependent covariates in the Cox and Fine-Gray models. Further-more, `FLORAL` is featured with built-in multi-step variable selection with further enhanced false discovery control and model interpretability (**Fig.1B**).

**Fig. 1:**
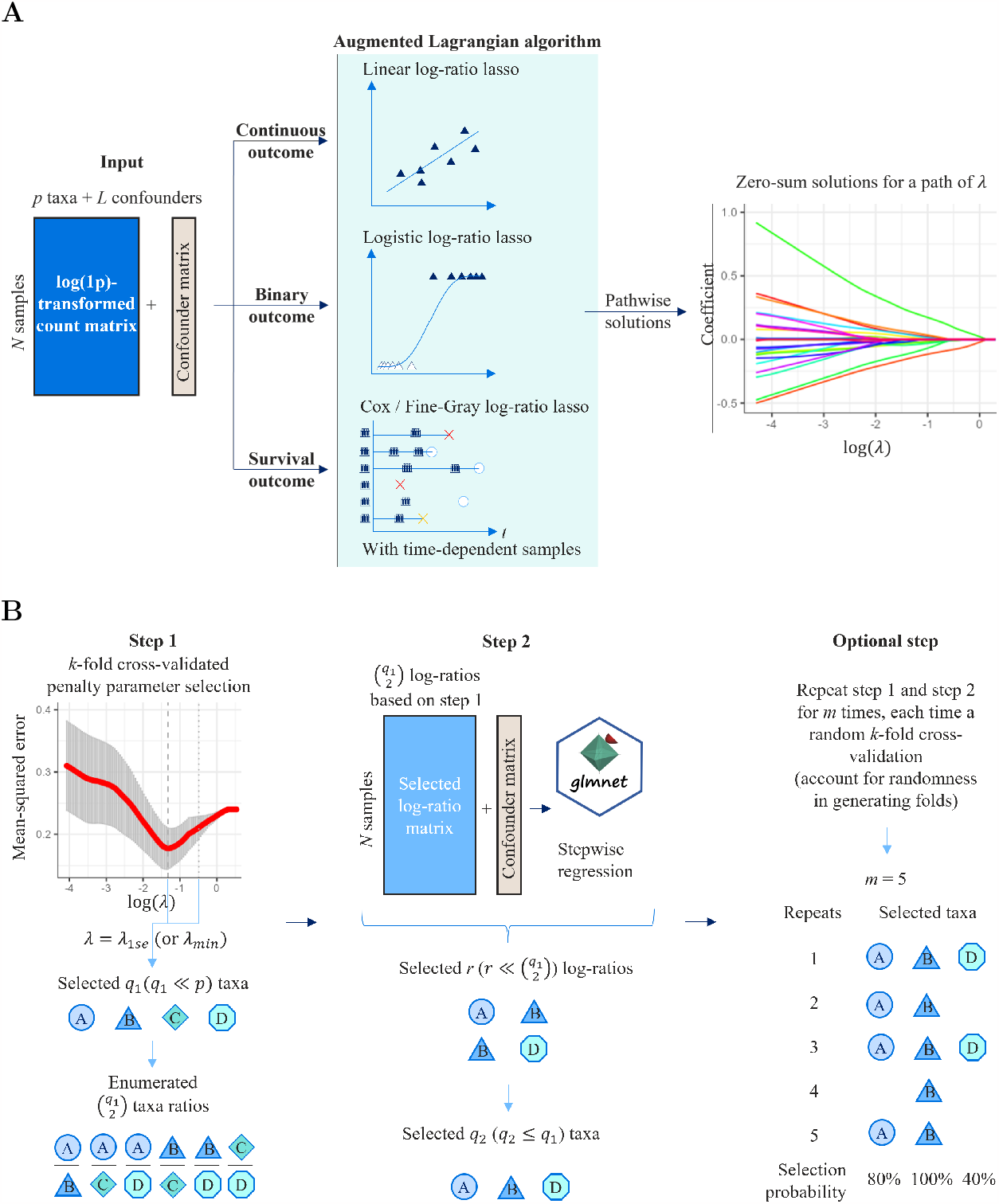
`FLORAL` performs log-ratio lasso regression and stepwise feature selection. **A**. The log-ratio lasso regression is conducted by an augmented Lagrangian algorithm with a zero-sum constraint, which is compatible with continuous, binary, survival and competing-risk outcomes. Longitudinal microbiome samples can be incorporated in survival models as time-dependent covariates. The algorithm is applied on a pre-specified path of penalty parameter *λ*, which returns a path of coefficient estimates satisfying the zero-sum constraint. **B**. Variable selection starts with *k*-fold cross validation (Step 1) which selects the penalty parameter and corresponding taxa with non-zero coefficients. The log-ratios enumerated from the taxa selected in Step 1 will be filtered in Step 2 by lasso regression and stepwise regression, where the remaining ratios and corresponding taxa are reported. Optionally, Step 1 and 2 can be repeated with additional *k*-fold data splits and calculations of taxa selection probabilities.

We conducted extensive real-data and simulation studies to assess our method’s performance and compare with various benchmark methods. We demonstrate that `FLORAL` achieves reasonable sensitivity and high specificity in publicly available microbiome datasets from 39 studies with binary comparison groups [27]. Using a 16S rRNA sequencing dataset of 8,967 longitudinal stool samples from a cohort of 1,415 allo-HCT patients from Memorial Sloan Kettering Cancer Center (MSKCC), we illustrate that incorporating longitudinal microbiome data can provide much richer information compared to only using baseline microbiome data, where we successfully identified *Enterococcus, Blautia, Erysipelatoclostridium*, and *Staphylococcus* as predictive features of patient overall survival, which have been previously reported [28–31].

## 2 Results

### 2.1 Simulations Showed Superior Variable Selection Performance of FLORAL Among Lasso-based Methods and Beyond

Extensive simulations based on the log-ratio models were performed to evaluate the sensitivity, specificity, and overall variable selection performance (*F*_1_ score) for different methods with various types of simulated outcomes, including continuous, binary, survival and competing-risks outcomes. Here, *F*_1_ score is defined as *F*_1_ = 2(precision^−1^+recall^−1^)^−1^ on a range between 0 and 1, where a higher *F*_1_ score indicates a better overall performance of precision and recall. We considered simulation scenarios with varying sample sizes (*n*), effect sizes (*u*), number of features (*p*), feature correlations (*ρ*), and feature sparsity levels (*s*), aiming to conduct a comprehensive method evaluation. In each simulation run, the outcome was generated based on log-ratios formed by 10 underlying “true” features. See Online Methods for detailed simulation configurations and descriptions of compared methods and evaluation metrics.

Our simulations demonstrated superior variable selection performance of `FLORAL` (**Fig.2, S1-S5**). As shown in **Fig.2**, `FLORAL` achieved the highest median *F*_1_ score out of 100 simulations in most scenarios with different sample sizes and types of outcomes, with big performance advantages for binary and survival outcomes under moderate sample sizes (*n* = 100, 200). Similar performance advantages were also observed under different effect sizes (**Fig.S2**), number of features (**Fig.S3**), correlation levels (**Fig.S4**) and sparsity levels (**Fig.S5**), with a few exceptions where `FLORAL` also reached comparable performances compared to other methods.

**Fig. 2:**
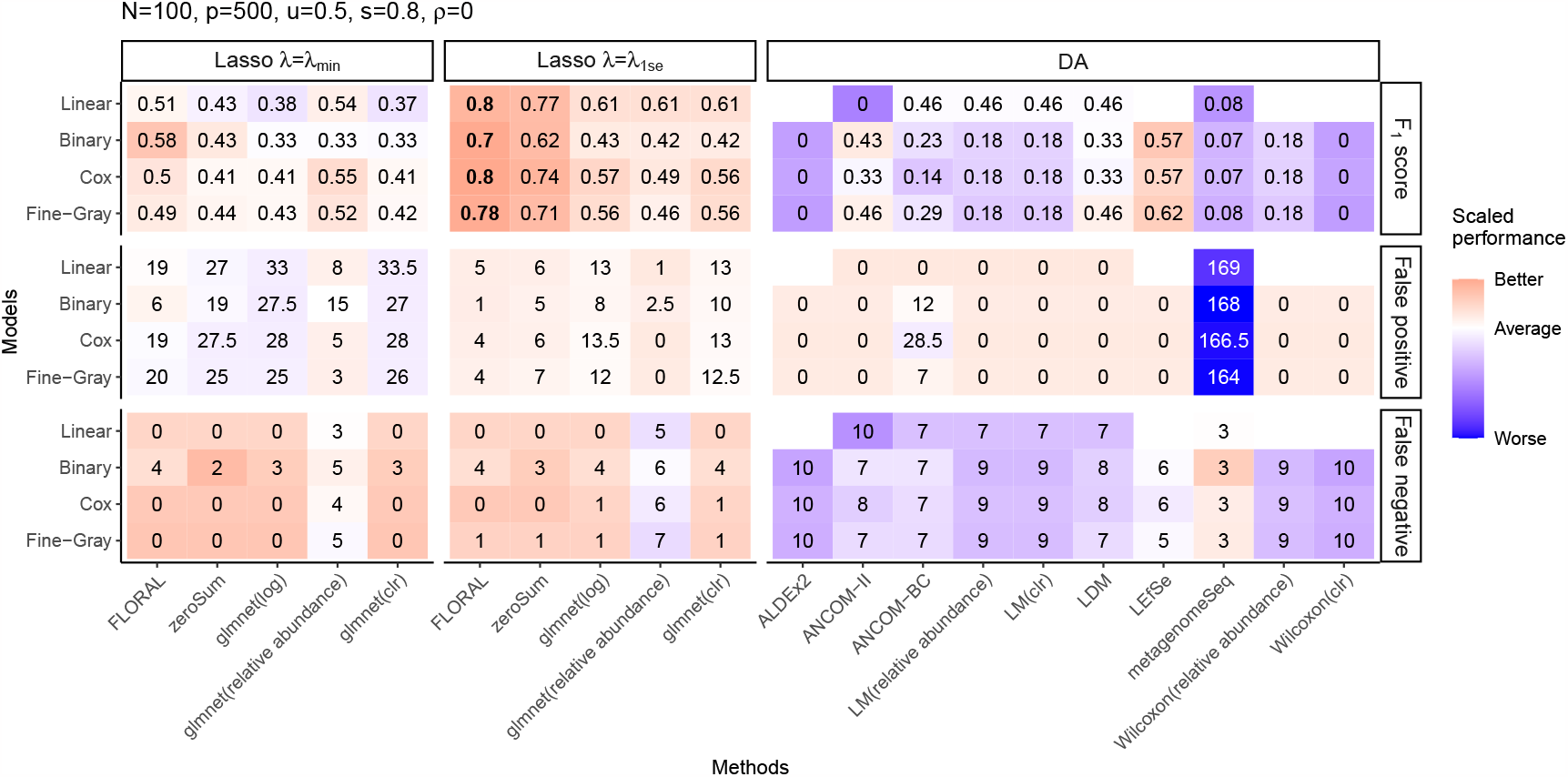
Median *F*_1_ score, median number of false positive features, and median number of false negative features obtained by lasso and DA methods for linear, binary, survival, and competing risk models out of 100 simulations with *n* = 100, *u* = 0.5, *p* = 500, *s* = 0.8, *ρ* = 0, where there were 10 true features out of *p* = 500 features in each simulation run. For each type of regression model, metrics across all methods were scaled to mean zero and standard deviation one for color visualization. The highest median *F*_1_ scores across all methods were printed in bold fonts. For the DA methods, the censoring indicator of the survival or competing risks outcomes were used to define patient groups except for LDM, where the Martingale residual was first computed then correlated with taxa abundances. Part of the DA methods were not evaluated for continuous outcome due to incompatibility. The adjusted p-value cutoff was set as 0.05 for all DA methods. log: log-transformation; clr: centered log-ratio transformation.

The better performance of `FLORAL` can be explained by the effective control of false positive features via its two-step feature selection mechanism (**Fig.S1A**) and the high sensitivity as an intrinsic characteristic of lasso-based method (**Fig.S1B**). Like other lasso-based methods, `FLORAL` obtained better overall *F*_1_ scores at *λ* = *λ*_1se_ than at *λ* = *λ*_min_ in most simulated scenarios, where a sparser selected feature set offered much fewer false positive features. Due to its stricter feature selection process, `FLORAL`’s sensitivity was slightly compromised when the effect size was very weak (*u* = 0.1, equivalent to odds ratio or hazard ratio of *e*^0.1^ = 1.1) or the sample size was small (*n* = 50) (**Fig.S1**,**S2**), where the setting *λ* = *λ*_1se_ could obtain zero selected features while *λ* = *λ*_min_ might reach a better *F*_1_ score. Nevertheless, `FLORAL` still achieved reasonable improvements over other lasso-based methods at fairly moderate effect sizes (*u* = 0.25, 0.5), larger sample sizes (*n ≥* 100) and various other settings.

Compared to `FLORAL` and other lasso-based methods, the DA methods showed generally lower false-positive rates but also much lower sensitivity at smaller sample sizes and moderate effect sizes (**Fig.S1B**,**S2C**), resulting in lower overall *F*_1_ scores (**Fig.2**,**S2A**). As sample size increased, the DA methods gained higher power to recognize true signals, gradually reaching or exceeding `FLORAL`’s *F*_1_ scores at sample size *n* = 500 (**Fig.2**). Notably, `metagenomeSeq` appeared to over-select features with a higher sensitivity but also much higher false-positive rates compared to other methods, while `ANCOM-BC` tended to have slightly inflated false positive rates at smaller sample sizes and smaller effect sizes (**Fig.S1A**,**S2B**). Moreover, `LDM` and `LEfSe` showed high robustness at reasonably large effect size (*u* = 0.5) and sample size (*n* = 200), where both methods maintained the best median *F*_1_ scores across all DA methods for binary and survival outcomes (**Fig.S3-S5**), outperforming `FLORAL` at smaller numbers of features (*p ≤* 200, **Fig.S3**) or at higher sparsity levels (*s* = 0.95, **Fig.S5**). This demonstrated the robustness of methods based on permutation test (`LDM`) and non-parametric test (`LEfSe`).

### 2.2 FLORAL Demonstrated Effective False Positive Control on 39 Publicly Available 16S rRNA Amplicon Sequencing Datasets

We applied various lasso-based regression methods and differential abundance testing methods on publicly available 16S microbiome datasets for 39 studies [32–67] as reported by Nearing et al. [27]. The 39 datasets contain a variety of research contexts including human, mouse, and environmental studies, where for each specific study there were two groups with hypothetical differences in their corresponding taxa abundance profiles. The distribution of sample size, number of features and the ratio between the comparison group sizes had a wide range, where both the sample size and number of features ranged from less than 50 to several thousands (**Fig.3A**). For lasso-based methods, we treated the identity of the binary comparison groups as a binary outcome, such that logistic regression with lasso penalty was performed.

**Fig. 3:**
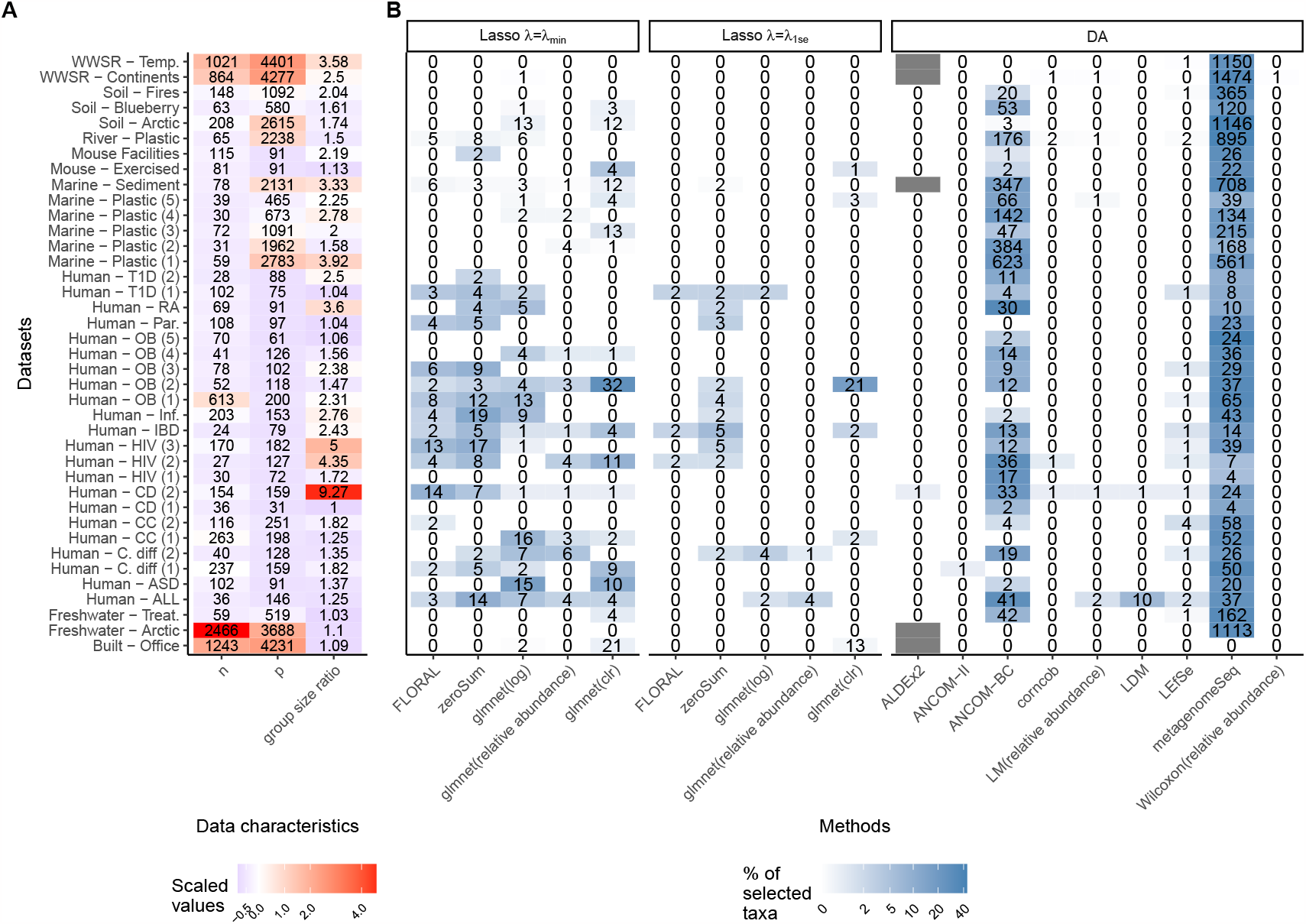
**A**. Data characteristics of the 39 publicly available 16S microbiome datasets, including sample size (n), number of genera (p), and ratio between the sizes of comparison groups. The color scheme represents scaled characteristics across all datasets. **B**. Number of selected taxa from the 39 publicly available 16S microbiome datasets by feature selection methods, with comparison group labels randomly shuffled. Part of data were unavailable for ALDEx2 due to memory overflow. The color scheme represents the percentage of selected taxa out of all taxa in a certain data set.

Due to the lack of gold standard definition of truly differentially abundant taxa, it is challenging to assess methods’ sensitivity. Therefore, we mainly focused on evaluating the specificity of the methods by randomly shuffling the group labels for each data set then running the methods. Theoretically, the differential abundance signals will be fully eliminated after random shuffling, such that any selected taxon can be treated as a false-positive feature. In parallel, we also applied the same methods on the original datasets without random group label shuffling, which offered descriptive statistics such as number of selected taxa, computing time, and median area under the receiver operating characteristic curve (AUC) of all selected taxa. See Methods section for detailed descriptions on the datasets and the configurations of different methods.

**Fig.3B** displays the numbers of selected taxa by various methods for the 39 public 16S datasets with shuffled group labels. As described above, larger numbers of selected taxa indicated higher false-positive rates. As observed, the lasso-based methods obtained reasonably low false-positive rates with the penalty parameter *λ* = *λ*_1se_, while there was an inflation of false positives when using *λ* = *λ*_min_. Thanks to its two-step variable selection strategy, `FLORAL` showed consistently lower numbers of false-positives than `zeroSum` while selecting zero taxa for all but three datasets with *λ* = *λ*_1se_. In terms of the DA methods, like observed in the simulations (Section 2.1), most methods selected zero taxa for most datasets and showed good false positive control. However, `metagenomeSeq` failed to control false positive findings, with false-positive rates up to 20% for most datasets. In addition, `ANCOM-BC` also had fairly high false-positive rates for datasets with relatively low sample sizes. The above observations were highly consistent with our simulation results (**Fig.S1A**), which further demonstrated `FLORAL`’s satisfactory protection against false positive findings.

The same analysis procedure was repeated for the original group labels without shuffling. The lasso-based approaches tended to select fewer genera than the DA methods (**Fig.S6**). This is expected as the DA methods perform comparisons for independent taxa then using multiple testing adjustment, such that many highly correlated features may be selected simultaneously. In contrast, lasso-based methods perform feature selection from multivariable regression models, such that the selected features are conditioned on all other feature values, resulting in a sparser set of selected taxa. Notably, `ANCOM-BC` and `metagenomeSeq` selected more genera than other methods for most datasets, which can be explained by their high false-positive rates as observed in **Fig.3B**. In addition, `FLORAL` achieved high median AUC (**Fig.S7**) and reasonable computing time (**Fig.S8**), showing good practical utility for datasets of diverse characteristics.

### 2.3 FLORAL Achieves Robust Signal Detection in Time-dependent Microbiome Samples

Allogeneic hematopoietic bone marrow transplant (allo-HCT) patients from Memorial Sloan Kettering Cancer Center (MSKCC) with eligible samples with 16S rRNA sequencing data between January 2009 and June 2021 were selected to investigate the associations between taxa abundance and patient overall survival (OS), transplant-related mortality (TRM) and graft versus host disease (GvHD)-related mortality (GRM). Here, TRM and GRM are defined as described by Copelan et al. [4] with relapse and progression of disease as competing risks. Two patient cohorts were derived, namely the peri-engraftment sample cohort and the longitudinal sample cohort (**Fig.S9**). The peri-engraftment sample cohort (912 patients, 912 samples) consisted of all patients with at least one sample collected between day 7 and 21 after transplant, where the latest collected sample was used as a peri-engraftment “baseline” sample, such that the microbial association with survival outcomes was investigated using only peri-engraftment samples. Accordingly, time to survival outcomes was landmarked at the sample collection day related to transplant. In contrast, the longitudinal sample cohort (1,415 patients, 8,967 samples) included all patients with samples available between day -30 to 730 relative to transplant, where the latest sample collected before the transplant day was regarded as the baseline (day 0) sample. Patients without available pre-transplant samples will enter the risk set of the survival models at days corresponding to their earliest available post-transplant samples. As listed in **Table S1**, patient characteristics of the two cohorts are largely similar, which created an ideal scenario to compare the strength of signals using peri-engraftment samples versus using longitudinal samples.

`FLORAL` was utilized to fit log-ratio lasso models with peri-engraftment samples and longitudinal samples for overall survival (Cox model), TRM (Fine-Gray model) and GRM (Fine-Gray model), where the penalty parameter was set as *λ* = *λ*_1se_ to enhance false-positive protection. In addition, the optional step of variable selection for drawing taxa selection probability was also applied, for 100 repeated 5-fold cross validations, to evaluate signal detection efficiency from either peri-engraftment samples or longitudinal samples. The regression models were adjusted for covariates including patient disease type, graft source, age, and conditioning intensity, where the lasso penalty was only applied to taxa features but not the covariates. We also applied other lasso-based methods and popular DA methods to investigate associations between genera and OS using the peri-engraftment and longitudinal sample cohort if compatible. See Online Methods for detailed description of the methods and cohorts used.

The taxa selection probabilities obtained from `FLORAL` demonstrated much stronger signal detection capability of longitudinal microbiome features compared to peri-engraftment microbiome features (**Fig.4**). Using the peri-engraftment sample cohort, the microbial feature detection rates were below 50% for all three considered survival endpoints, indicating feature detection was largely dependent on the fold split and was less reliable (**Fig.4A-C**). In contrast, The longitudinal sample cohort provided not only more samples per patient but also more patients with eligible samples, which helped identify genera with detection rates higher than 80% or even 100% (**Fig.4D-F**). In particular, genera *Enterococcus, Blautia, Erysipelatoclostridium* and *Staphylococcus* were selected from the longitudinal sample cohort with high selection probabilities. Specifically, *Enterococcus* and *Staphylococcus* showed consistently harmful associations with OS, TRM and GRM, *Blautia* were identified to be associated with better OS and lower GRM cumulative incidence, and *Erysipelatoclostridium* were found to be also associated with better OS, and lower TRM and GRM cumulative incidences. Such high selection probability for the above three genera was not seen from the models using the peri-engraftment sample cohort (**Fig.4A-C**). The above results from the longitudinal sample cohort were highly consistent with previous studies [28–31, 68], demonstrating powerful utilities of `FLORAL` in analyzing survival endpoints with longitudinal microbiome data, where the signal detection is much more robust than using a single-time microbiome sample for each patient.

**Fig. 4:**
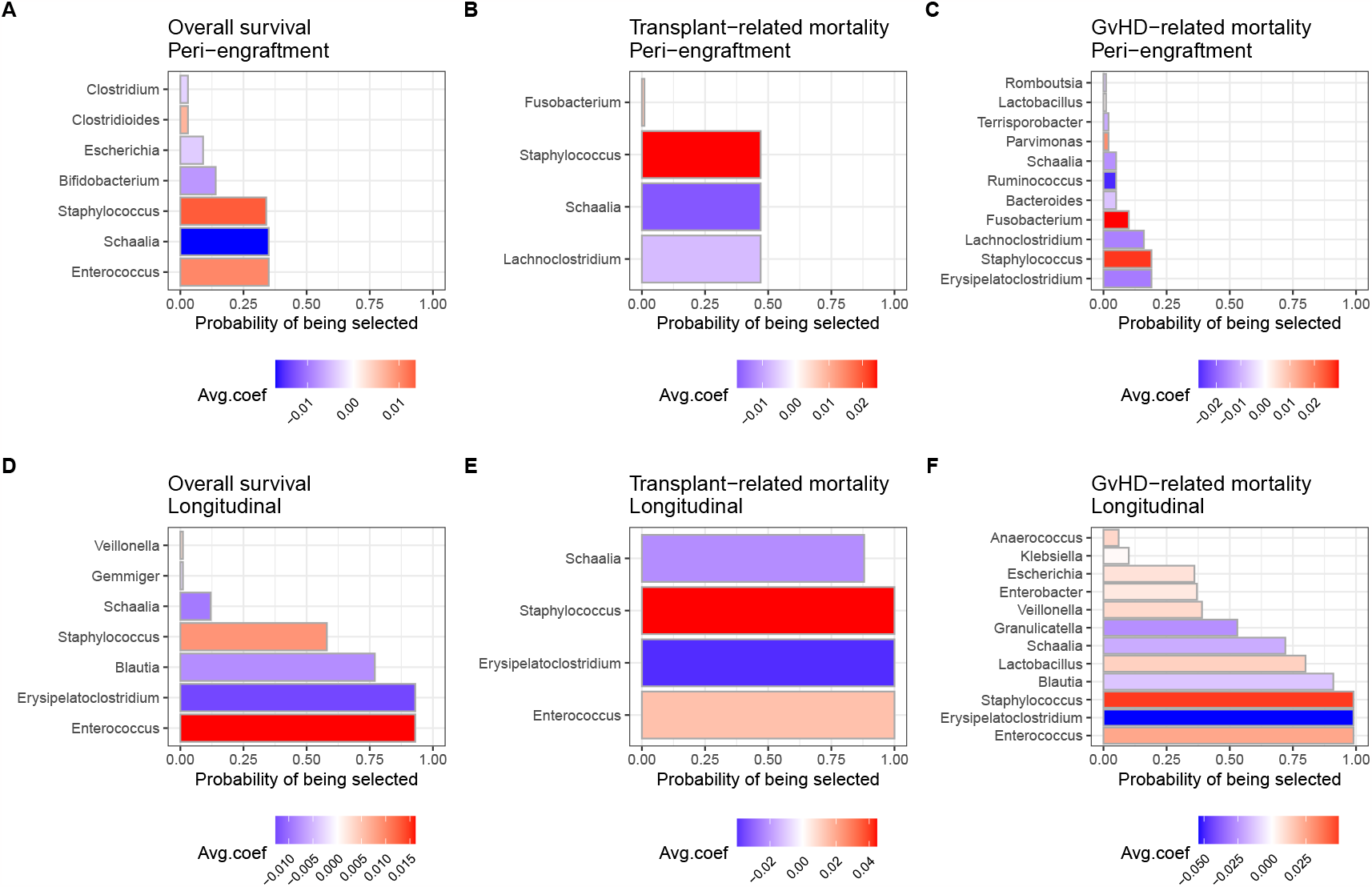
Probabilities of genera being selected from 100 repeats of 5-fold cross-validation with random fold split for Cox model of overall survival with **A**. peri-engraftment samples or **D**. longitudinal samples; Fine-Gray model of transplant-related mortality with **B**. peri-engraftment samples or **E**. longitudinal samples; Fine-Gray model of GvHD-related mortality with **C**. peri-engraftment samples or **F**. longitudinal samples. The color scheme represents the average lasso coefficient estimates of the corresponding genus at *λ*^(*i*)^ = *λ*_1se_ over *i* = 1, …, 100 repeats.

Compared to `FLORAL`, other lasso-based methods and popular DA methods did not achieve as effective feature selection performances. Like `FLORAL`, `glmnet`-based lasso models can also incorporate longitudinal microbial features with different data transformation options. However, these methods reached much lower feature selection rates than `FLORAL` using the longitudinal sample cohort in 100 cross-validation runs (**Fig.S10**), where important genera such as *Enterococcus* were hardly detected. In addition, `zeroSum` and `glmnet` were not able to better detect important genera using the peri-engraftment sample cohort than `FLORAL` (**Fig.S11**), indicating weak signals when only using the peri-engraftment microbiome samples. Unlike `FLORAL` and `glmnet`, the DA methods are incompatible with longitudinal microbiome samples, and thus were only applied for the peri-engraftment cohort. As shown in **Fig.S12**, many DA methods conservatively selected no features at the threshold of 0.05 for the adjusted p-value, while `LEfSe` and `metagenomeSeq` selected a large number of genera. Nevertheless, all DA methods failed to identify *Blautia* and *Erysipelatoclostridium* as detected by `FLORAL` using the longitudinal sample cohort. The above results suggest that `FLORAL`’s improvements in microbial feature selection from the peri-engraftment cohort to the longitudinal cohort are attributed not only to its flexibility of incorporating longitudinal microbial features as time-dependent covariates, but also to its infrastructure of utilizing log-ratio based regression models.

## 3 Discussion

In this work, we present `FLORAL` for fitting log-ratio lasso regression models powered by the augmented Lagrangian algorithm with a two-step variable selection procedure. Compared to existing log-ratio lasso methods, `FLORAL` maintains reasonable sensitivity in variable selection, shows better false positive control in real data analyses, and effectively improves signal detection by incorporating longitudinal microbial features as time-dependent covariates.

Compared to the widely applied microbiome data transformation based on relative abundance 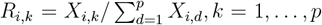, the log-ratio model better fits the compositional nature of microbiome data and provides several conveniences in handling the data and interpreting microbial associations. First, relative abundance *R*_*i,k*_ depends on the absolute counts of the collection of *p* taxa measured from the sequencing process. Given different sequencing depths across samples or studies, the detectable taxa features vary, which may affect the consistency in quantifying *R*_*i,k*_. This challenge was earlier described as “subcomposition difficulty” [19], which leads to different analysis results due to the varying definition of the entire feature set. In contrast, the ratio *X*_*i,j*_*/X*_*i,k*_ between two taxa *j* and *k* is a stable quantity that is invariant of subcomposition changes across samples caused by technical variations, which can potentially enhance the reproducibility of analyses. Moreover, taxa-specific bias is highly prevalent in microbiome data [18], such that only ratios between two taxa carry stable relative magnitudes across samples or studies that is invariant to taxa-specific biases, which further supports the analysis based on ratios over relative abundance.

As demonstrated by simulation and real data studies, the two choices of the penalty parameter *λ*_min_ and *λ*_1se_ have different properties, where *λ*_1se_ achieved better *F*_1_ scores in simulations and lower false positive rates in the analyses of 39 real datasets compared to *λ*_min_. Therefore, we recommend users to choose *λ* = *λ*_1se_ for better control of false discoveries. In studies with smaller sample sizes, it is likely to detect zero features with *λ* = *λ*_1se_ in a single two-step variable selection with cross-validation. For those studies with small scales, we recommend using multiple runs of cross-validation to rank the strength of the features by their selection probabilities, such that features with weak signals may still be captured and reported regarding their importance relative to other features.

The proposed method offers an effective alternative to the popular differential abundance testing approaches. Unlike the DA approaches where users are required to specify cutoffs for adjusted p-values, `FLORAL` conducts variable selection based on cross-validated prediction error or model fitting criteria, such that the selected taxa have direct contribution to better prediction performances and are not determined by arbitrary p-value thresholds. Moreover, the log-ratio lasso regression method better addresses the association between survival outcomes and microbiome data and offers a natural framework for incorporating longitudinal microbial features, which appeared to be challenging for the DA methods. However, it is important to note that the DA methods may serve as more reasonable options if the research interest is to compare paired or correlated microbial features [12], where generalized estimating equation (GEE) extensions of log-ratio lasso regression are required to better account for the dependency across subjects.

In large-scale follow up studies with longitudinal samples, one commonly encountered challenge is to utilize all of the microbiome data. Due to the limitation of the DA methods, it is usually only possible to perform two-sample comparisons for microbiome samples collected at a specified time window, such as the peri-engraftment period in the allo-HCT patient cohort. Although linear mixed-effect models (`MaAsLin2`, for example) have been proposed for longitudinal microbiome data analysis [2], the method can be regarded as an extended DA method in the sense that it requires a pre-specified significance threshold, clearly defined groups for comparison, and a data transformation scheme which is usually based on relative abundance. When the comparison groups are well defined at baseline, such as treatment group versus control group, it is helpful to apply linear mixed-effect models to investigate the association between the comparison groups and microbial trajectories. On the other hand, if a survival endpoint is of interest, then a regression model with time-dependent covariates, like `FLORAL`, will be more appropriate to better incorporate time-to-event information.

There are several opportunities of further development for `FLORAL`. First, the regularization model can be extended beyond the scope of the lasso regression with *ℓ*_1_-penalty, which facilitates subsequent fine-tuning of the models with potential utility of prediction. Our augmented Lagrangian algorithm can be easily modified to perform elastic-net regression [69], adaptive lasso [70], or other regularization forms. Second, it is of high interest for medical researchers to perform inference on selected features, which motivates developments of post-selection inference procedures for the log-ratio lasso models. Last but not least, the application of `FLORAL` can also be extended to other compositional biomedical data, such as cell ratios from flow cytometry or single-cell sequencing experiments, nutrient ratios from dietary data, and metabolomics data.

## Acknowledgments

This research was supported by National Cancer Institute award numbers, R01-CA228358, R01-CA228308, P30 CA008748 MSK Cancer Center Support Grant/Core Grant and P01-CA023766; National Heart, Lung, and Blood Institute (NHLBI) award number R01-HL123340 and R01-HL147584; National Institute on Aging award number P01-AG052359, and Tri-Institutional Stem Cell Initiative. Additional funding was received from The Lymphoma Foundation, The Susan and Peter Solomon Family Fund, The Solomon Microbiome Nutrition and Cancer Program, Cycle for Survival, Parker Institute for Cancer Immunotherapy, Paula and Rodger Riney Multiple Myeloma Research Initiative, Starr Cancer Consortium, and Seres Therapeutics. OM reports funding from the Hyundai Hope on Wheels Young Investigator Award and Tow Center for Developmental Oncology Career Development Award. ML reports funding from Beijing Nova Program of Science and Technology (Z191100001119120). JUP reports funding from NHLBI NIH Award K08HL143189 and the V Foundation.

## Methods

### 4.1 Overview of FLORAL

Given *p* microbial features, *L* confounding factors (if applicable), and the corresponding outcome of interest, `FLORAL` performs log-ratio lasso regression and subsequent variable selection for continuous, binary, and survival or competing risk outcomes (**Fig.1**), where longitudinal microbial features are incorporated in time-to-event models as time-dependent covariates. The regression model assumes that only a sparse set of the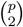 possible ratios between two microbial features are associated with the outcome of interest, which can be achieved by *ℓ*_1_-regularization on a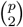-dimensional unknown parameter space. An augmented Lagrangian algorithm with a zero-sum constraint, which effectively reduces the covariate space from 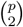dimensions to *p* dimensions, was developed to conduct pathwise estimation of the log-ratio lasso models under pre-specified values of the penalty parameter *λ*. Subsequently, the step 1 variable selection is based on a *k*-fold cross validation, where a cross-validated predictive model assessment metric (such as mean-squared error or deviance) helps to identify a value of *λ* which achieves the best prediction (*λ*_min_) or a sparser feature set with reasonable model fitting (*λ*_1se_). Given selected *λ*, the *q*_1_ taxa (*q*_1_ *<< p*) with non-zero regression coefficients will be selected as taxa contributing to better prediction performances. With *q*_1_ selected taxa, `FLORAL` enumerates all 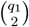 possible ratio configurations, then performs the step 2 variable selection by running lasso regression followed by stepwise regression on the 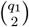-dimensional log-ratio features, which further selects *r* ratios 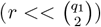 with strongest signals. Subsequently, *q*_2_ taxa (*q*_2_ *≤ q*_1_) forming the *r* selected ratios can be obtained as selected set of predictive taxa. As an optional step, variable selection steps 1 and 2 can be repeated for *m* times, such that the variable selection can be replicated under multiple random configurations of folds for cross validations. This optional step can help assess the probability of a certain taxon being selected after accounting for the uncertainties in defining folds.

### 4.2 Log-ratio Regression Models

For a given sample, let ***X*** denote the absolute count vector for *p* microbial taxa. Let ***W*** denote the confounder vector with *L* features. For a scalar outcome such as continuous or binary outcome, we denote the corresponding response variable as *Y*. For survival outcome, we denote 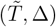 as observed survival time subject to right censoring and the censoring indicator, respectively. We denote the realization of the above random quantities for the *i*th patient as 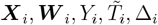, respectively. For a scalar outcome *Y*_*i*_, we model the association between *Y*_*i*_ and ***X***_*i*_, ***W*** _*i*_ via a log-ratio generalized linear regression model (GLM):

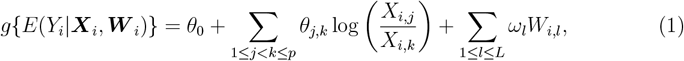

where *g*(·) is a link function accounting for the distribution of ***Y***, *θ*_0_ is an unknown intercept term, *A*_*i,j*_ represents the *j*th element of the vector ***A***_*i*_, and *θ*_*j,k*_, *ω*_*l*_ are unknown coefficients corresponding to the paired log-ratios log(*X*_*i,j*_*/X*_*i,k*_) and the patient characteristics *W*_*i,l*_, respectively. Here, we adapt the notion of pairwise log-ratio [22], where there are 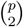 unknown *θ*_*j,k*_ for 1 *≤ j < k ≤ p*. Similarly, for survival outcome 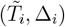, we consider a log-ratio proportional hazards model

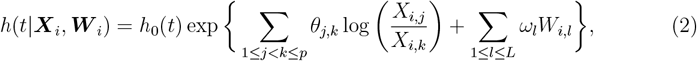

where *h*(*t*|***X***_*i*_, ***W*** _*i*_) denotes the hazard function conditioned on microbial features ***X***_*i*_ and patient characteristics ***W*** _*i*_, and *h*_0_(*t*) is the baseline hazard function. Note that model (2) can be naturally extended for longitudinal microbiome data ***X***(*t*), such that

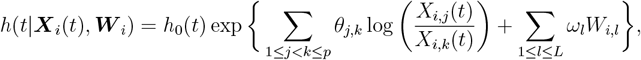

where ***X***(*t*) can be updated at different times of sample collection. In practice, longitudinal microbiome samples are only available at a finite number of time points, where the last value carried forward (LVCF) strategy is applied [71]. Moreover, the Fine-Gray subdistributional proportional hazards model [26] can be equivalently estimated by a weighted Cox model [72], which offers a convenient pathway of implementing competing risks modeling under the same framework.

Both models (1) and (2) can be simplified as a more concise form by rewriting the log of ratios as differences of log-counts [22, 23]. Let 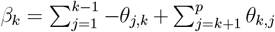, one can show by algebra that

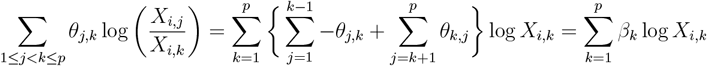

and

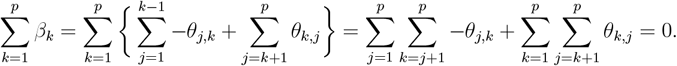

Therefore, models (1) and (2) can be rewritten as

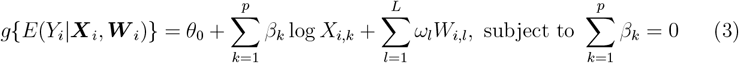

and

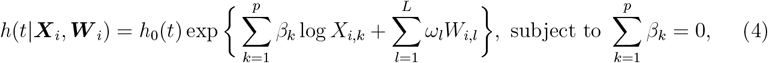

correspondingly. In modern microbiome studies, the number of taxa *p* can reach the scale of thousands. Compared to models (1) and (2) which impose 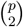 log-ratio features, models (3) and (4) show appealing computational benefits of having a much lower dimensional covariate space as *p* increases. To address commonly encountered zero counts in microbiome data, we suggest using log(***X*** + 1) to replace log(***X***) as an approximate covariate space which keeps zero counts as zeros after log transformation.

### 4.3 The Log-ratio Lasso Estimator and the Augmented Lagrangian Algorithm

Denote ***β*** = (*β*_1_, …, *β*_*p*_)^*T*^, ***ω*** = (*ω*_1_, …, *ω*_*L*_)^*T*^, and ***ζ*** = (*θ*_0_, ***β***^*T*^, ***ω***^*T*^)^*T*^ for the GLM model or ***ζ*** = (***β***^*T*^, ***ω***^*T*^)^*T*^ for the proportional hazards model. Let *L*(***ζ***) denote the log-likelihood of model (3) or the log-partial likelihood of model (4). We define the log-ratio lasso estimator as

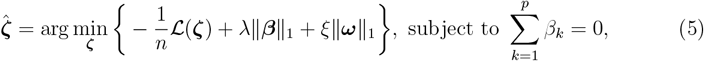

where *λ* and *ξ* are regularization penalty parameters and ∥ *·* ∥_1_ denotes *ℓ*_1_-norm. Here, we consider different regularization parameters for ***β*** and ***ω*** to facilitate higher flexibility in real practice, where investigators may set *ξ* = *λ* or *ξ* = 0 to conduct microbial feature selections with or without penalizing the confounding covariate effects.

We adapt the similar treatment in `glmnet` [73] to approximate *ℒ*(***ζ***) by its second-order Taylor expansion centered at 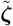, which is either a vector of initial values or the estimates from a previous iteration. Let 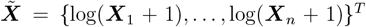 and 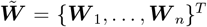 denote the *n × p* log-transformed microbiome count matrix and the *n× L* confounding covariate matrix, respectively. Define 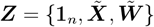 (GLM) or 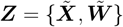(proportional hazards model), 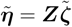 as the linear predictor with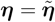, and 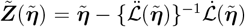, where 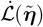and 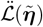 denote the gradient and Hessian matrix of 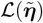, respectively. Then by second-order Taylor expansion we have

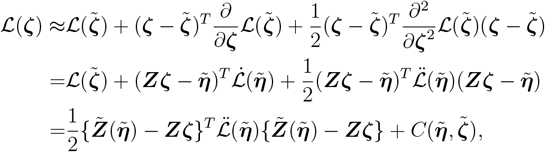

where the first term in the formula on the last row is a weighted quadratic form of ***Zζ*** and the second term 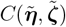 is independent of ***ζ***. To alleviate computational burdens for the *n×n* matrix 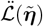, we follow [69] and [74] to substitute 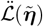 by its diagonal elements 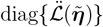. That is, the working loss function is defined as

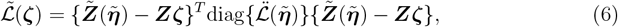

which is a standard weighted least squares form with continuous response vector 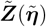 and predictor matrix ***Z***. It is also straightforward to show that 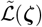 is equivalent to the standard least squares form 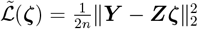 when ***Y*** is continuous. Based on the working loss function, we obtain the working optimization problem for the proposed lasso estimator:

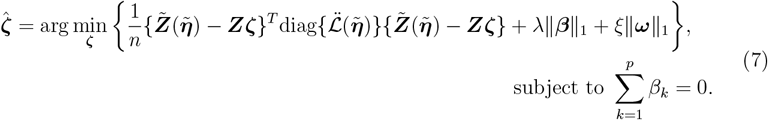

With a unified formula of working loss function (6) for different choices of *L*(***ζ***), the corresponding lasso optimization problem with constraint (7) can be conveniently defined for either scalar or survival outcomes if the first- and second-order differentiation with respect to 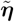 are well defined for the log-likelihood, or partial log-likelihood function *ℒ*(***ζ***).

We adapted the augmented Lagrangian approach [75] to solve the constrained optimization problem (7). Specifically, the constraint 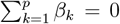 is incorporated in the following target function

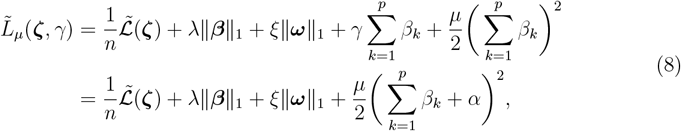

where *γ* is the Lagrange multiplier, 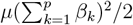 is the standard term used in the penalty method. Following Lin et al.’s approach [20], we define *α* = *γ/µ* enables merging the Lagrange multiplier 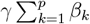 and the penalty term 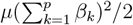 into a single term. In practice, the augmented Lagrangian method is able to achieve the constraint without using a overly large value of *µ*, which avoids ill-conditioning caused by having large *µ* [76]. We typically let *µ* = 1 as fixed in our algorithm.

Given *λ, ξ, µ*, and an initial value 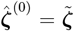 which can be obtained by a warm start, estimation of 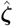 is conducted by a coordinate gradient descent algorithm with iteratively updated value of *α*, where the initial value of *α* at the first iteration, *α*^(0)^, is zero. In the *i*th iteration, the corresponding estimate 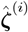 is updated by minimizing 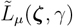 with fixed values of *λ, ξ, µ, α*^(*i*)^ and an initial value 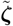 from the previous iteration. This step of updating 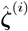 can be performed by an inner loop of standard coordinate descent algorithm. Specifically, if the *k*th component of 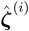 is the *h*th component of 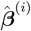, then it will be updated by

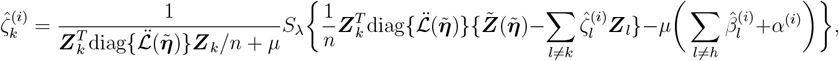

where ***Z***_*k*_ denotes the *k*th column of matrix ***Z*** and *S*_*λ*_(*x*) = sgn(|*x*| − *λ*)_+_ is the soft thresholding operator. Similarly, if the *m*th component of 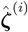 belongs to 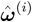, then it is updated by

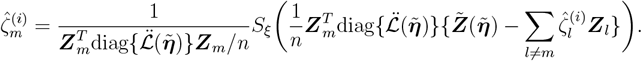

As observed from the above update formula for 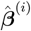 and 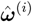, the main difference is that the zero-sum constraint is only applied for 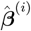, but not for 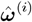. After each inner-loop coordinate descent for each feature, we update 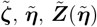, and 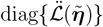 for the next inner-loop coordinate descent. The updates of 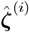 stops if the loss function 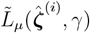 is converged at a tolerance parameter *δ*′. With updated 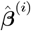 from the inner loop, the penalty parameter *α* is updated as

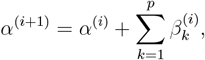

such that a larger penalty will be imposed in the (*i* + 1)th iteration for 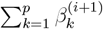 if 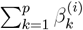 deviates from zero in the *i*th iteration. The algorithm stops if ∥***ζ***^(*i*)^−***ζ***^(*i*−1)^∥_1_ *< δ* for a pre-specified tolerance parameter *δ*. Detailed implementation of the algorithm is reported as Algorithm 1. In actual implementation, we calculate *p × p* matrix ***A*** and *p*-vector ***B***, as defined in Algorithm 1, prior to the coordinate descent loop to save computational cost. We also specify the maximum iteration number *u*^*′*^ for the inner loop and *u* for the outer loop to bring an early stop if the convergence is not reached. In our analysis, we constantly use *µ* = 1, *δ* = *δ*^*′*^ = 10^−7^ and *u* = *u*^*′*^ = 100.

### 2.4 Pathwise Solution and Cross Validation

To have a global picture on how feature sparsity is governed by different choices of *λ*, we solve the optimization problem (7) by Algorithm 1 on a decreasing path ***λ*** of *λ*. By default, the path ***λ*** starts with

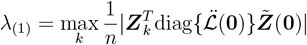

which acquires 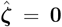 [74]. Then a sequence of length *m, λ*_(1)_, …, *λ*_(*m*)_ is generated with equal distance on log scale, where *λ*_(*m*)_ is typically selected as 0.01*λ*_(1)_ if *n < p* and 0.0001*λ*_(1)_ if *n ≥ p*. Here we consider *ξ* = *λ* or *ξ* = 0, such that *ξ* follows the same path as *λ* does or is fixed as a constant.

*k*-fold cross validation is used to determine the optimal choice of *λ* which maximizes the cross-validated predictive performance. Standard criteria, such as mean-squared error and deviance are used to evaluate prediction errors. Two choices of *λ* are reported, namely *λ*_min_ which minimizes cross-validated prediction error, and *λ*_1se_ which provides a sparser solution than *λ*_min_ but still obtains cross-validated prediction error within one standard error of that of *λ*_min_. The cross validation serves as the first step of variable selection in `FLORAL` (**Fig.1B**), where taxa with non-zero coefficient estimates 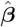 at *λ*_min_ or *λ*_1se_ are selected for step 2 variable selection.

### 4.5 Step 2 Variable Selection

In the previous sections we derive the algorithm to efficiently find a sparse set of predictive taxa. However, specific pairs of log-ratios are not identifiable via cross validation based on the estimates obtained by Algorithm 1. To facilitate a sparser feature selection with interpretability for specific ratios, one natural extension is to perform exhaustive search on all possible pairs of the log-ratios for the selected features obtained from the Step 1 cross validation [22]. Since the number of selected taxa *q*_1_ from the Step 1 cross validation is much smaller than *p*, the corresponding number of pairwise combinations 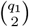 is also much smaller than 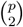 (**Fig.1B**), which only requires standard memory usage in standard R packages for lasso models and stepwise regression. In our implementation, we first perform a standard lasso regression via `glmnet` over the enumeration of log-ratios from the selected feature set to filter out log-ratios not contributing to a better prediction. Then we apply a stepwise regression model for the selected log-ratios to further exclude log-ratios that do not substantially improve model fitting. The two-stage feature selection aims to keep the strongest signals in the model while obtaining meaningful interpretations for specific ratios of microbes.

### 4.6 An Optional Step for Feature Selection Probabilities

The result of the cross-validated variable selection and the subsequent second step selection depends on how subjects are split into folds, such that different fold splits may select different taxa or taxa ratios. In real data analysis where sample size is small or signals are weak, it is helpful to repeat the cross validation for more objective evaluations of feature selection.

Thus, we developed an optional step to assess the reliability of variable selection, which repeats the *k*-fold cross-validated 2-step variable selection procedure by *m* times (**Fig.1B**). In each of the *m* repeats, the cross validation folds will be randomly generated, such that the corresponding penalty parameter *λ* will correspond to different sets of selected features. This optional step allows investigators to assess how robustly a certain microbial feature is selected based on different fold split schemes, where a higher selection probability indicates higher confidence of association between the feature and the outcome.

### 4.7 Simulation Studies

#### 4.7.1 Data Generation

We performed extensive simulation studies to assess various methods’ performances under different scenarios. Let *n* be the sample size and *p* be the number of features. For each simulated sample *i, i* = 1, …, *n*, we first simulate the underlying taxa composition ***c***_*i*_ = (*c*_*i*1_, …, *c*_*ip*_)^*T*^, where *c*_*ik*_ *≥* 0 for all *k* and 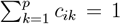. To get ***c***_*i*_, we simulate a *p*-vector ***x***_*i*_ which follows a *p*-variate normal distribution *N*_*p*_(***ξ*, Σ**), where ***ξ***_*k*_ = log *p* for *k* = 1, 2, 3, 5, 6, 8 and otherwise ***ξ***_*k*_ = 0. This choice of ***ξ*** makes features 1, 2, 3, 5, 6, 8 more abundant than others. Correspondingly, the variance parameters 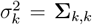 satisfies 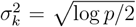 for *k* = 1, 2, 3, 5, 6, 8 and otherwise 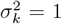, which makes highly abundant features of higher variation. We let Σ_*j,k*_ = *ρ*^|*j*−*k*|^, *ρ ∈* (0, 1) be the correlation between features *j* and *k*, where features of adjacent indices are more correlated than features of distant indices. To generate sparsity in counts, we then specify a sparsity level *s ∈* (0, 1) and randomly force *s × p* many elements in ***x***_*i*_ to be −*∞*. Then we calculate *c*_*ik*_ = exp(***x***_*ik*_)*/*∑_*d*_ exp(***x***_*id*_) for all *k* to obtain ***c***_*i*_.

Four types of outcomes, namely continuous, binary, time-to-event, and competing risk, are considered in our simulations conditioned on ***c***_*i*_. Given ***c***_*i*_, we first generate a “true count” vector ***C***_*i*_ following a multinomial distribution with 10^6^ counts and probability vector ***c***_*i*_. Note the ***C***_*i*_ facilitates defining the log-ratios by log(1+ ·) transformation, which mitigates the arbitrary choice of increments for proportions ***c***_*i*_. Then the corresponding underlying true linear predictor is generated as

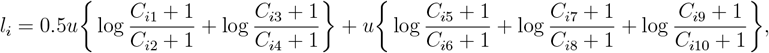

such that the first ten simulated features are true features associated with the outcome in the form of log-ratios. Here *u* controls the effect sizes, where the first two ratios have half of the effect sizes of the latter three ratios. Given *l*_*i*_, the continuous response variable 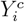 is generated by

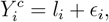

where the error term *ϵ*_*i*_ follows independent standard normal distribution for each *i*. Similarly, the binary outcome variable 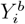 is simulated from a *Bernoulli*(*q*_*i*_) distribution, where *q*_*i*_ = expit(*l*_*i*_) and expit(*x*) = 1*/*(1+*e*^−*x*^). For the time-to-event outcome,, the event time *T*_*i*_ is simulated from the distribution function 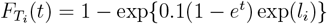.

It can be shown that the associated hazard function is equal to 0.1*e*^*t*^ exp(*l*_*i*_) which belongs to the family of proportional hazards models. Then a random censoring time *V*_*i*_ is generated as the minimum of an *Exponential*(0.1) and a *Uniform*(5, 6) distribution. Then the observable survival time 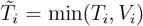 and event indicator Δ_*i*_ = *I*(*T*_*i*_ *< V*_*i*_) are obtained. For the competing risk outcomes, we follow Scheike et al.’s simulation approach [77]. Specifically, two failure types are assumed, where the cumulative incidence of the first and second failure types satisfy 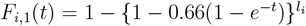 and 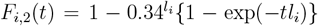, respectively. The failure type *ϵ*_*i*_ ∈ {1, 2} can then be generated by the failure type probabilities defined by *F*_*i*,1_(*∞*) and *F*_*i*,2_(*∞*). Given failure type *ϵ*_*i*_, the failure time *T*_*i*_ is generated from the conditional distribution function 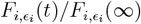. An Independent censoring time *V*_*i*_ is independently generated from a *Unif* (0.19, 10). Then the observable survival time 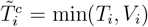 and failure type indicator 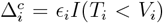 are obtained. In data analysis, we focus on investigating the association between features and the first of the two failure types.

Based on the underlying true taxa composition ***c***_*i*_, we further simulate the observable count data ***X***_*i*_ with different sequencing depths. First, the sequencing depth *D*_*i*_ is generated as the largest integer smaller than a random variable following a *Unif* (5000, 50000) distribution, where 5000 to 50000 is a reasonable range for high-quality microbiome 16S rRNA sequencing depths. Then the count data ***X***_*i*_ is generated from a multinomial distribution with *D*_*i*_ instances and the probability vector ***c***_*i*_.

For each of the four types of outcome variables, we investigated the performance of methods based on a reference scenario where *n* = 200, *p* = 500, *s* = 0.8, *ρ* = 0, and *u* = 0.5. Controlling other parameters as fixed, we compared *n* = 50, 100, 200, 500, *p* = 100, 200, 500, 1000, *s* = 0.8, 0.95, *ρ* = 0, 0.5, and *u* = 0.1, 0.25, 0.5. This serves as a comprehensive survey in understanding the behavior of methods under various settings. For each simulation run, the simulated outcome 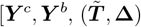, or 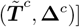 and observable count matrix 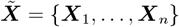 were the input data to various methods.

#### 4.7.2 Method Configuration and Assessment

We tested lasso-based methods and different abundance (DA) testing methods in simulations as listed in **Fig.2**. For lasso-based methods, we considered methods with zero-sum constrained lasso (`FLORAL` and `zeroSum`) and standard `glmnet` models with relative abundance, centered log-ratio transformed counts, and log-transformed counts. The same random fold split was used for all methods, where 10-fold cross-validated mean-squared error was used to identify *λ*_min_ and *λ*_1se_ for scalar outcomes *Y* ^*c*^ and *Y* ^*b*^, while 10-fold cross-validated log-likelihood deviance was used for survival outcomes 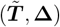 and 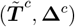.

Features with non-zero coefficients at chosen values of penalty parameter were regarded as selected features. For `FLORAL`, remaining features after the Step 2 variable selection were used for method assessment.

For the DA methods, we largely applied the methods with their default configurations as detailed in **Table S2**. In addition, the Benjamini-Hochberg approach was applied for p-value adjustments if applicable, where taxa with adjusted p-values smaller than 0.05 were defined as selected features. Scalar outcomes ***Y*** ^*c*^ or ***Y*** ^*b*^ were treated as covariates for the DA methods. We did not test `ALDEx2`, `LEfSe`, nor the Wilcoxon test for continuous outcomes ***Y*** ^*c*^ due to incompatibility. For time-to-event outcome 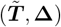 or 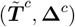, we used the Martingale residual as the covariate for `LDM`, while the censoring indicator **Δ** or *I*(**Δ**^*c*^ = 1) was used as the patient group indicator for other methods. Detailed versions of R packages used for each method are listed in **Table S2**.

We focused on evaluating the variable selection performance for each method. Given the knowledge of the ten underlying truly associated features, we summarized the number of false negatives (FN, ranging between 0 and 10), false positives (FP, ranging between 0 and *p* − 10), and the *F*_1_ score 2TP*/*(2TP + FP + FN) (ranging between 0 and 1), for each method at each simulation run, where TP represents the number of true positives. Here, a smaller FN indicates better sensitivity, while a smaller FP indicates better specificity of the methods. Similarly, a higher *F*_1_ score implies a better balance between precision and recall. To better visualize the simulation results, heatmaps for median *F*_1_, median FN, and median FP were generated with colors scaled for each simulation scenario. Simulation results from `corncob` was omitted in the figures as we observed zero features being selected in all simulation scenarios, which implies that the data generating model does not satisfy `corncob`’s model assumption.

### 4.8 Real Data Applications

#### 4.8.1 Publicly Available Datasets for Two-Sample Comparison

Publicly available 16S rRNA sequencing datasets from 39 studies [32–67] were retrieved from the online data repository [78]. We applied the same naming system as used by Nearing et al. [27] to annotate the datasets, which are presented in **Fig.3A**. For each dataset, sequencing counts from different amplicon sequencing variants (ASVs) but the same genus were aggregated to form the genera count table for subsequent analysis. All counts were included without pre-filtering. Appropriate transformations were applied if applicable to obtain data formats suitable for different methods. For linear regression model (LM) and Wilcoxon test, we used relative abundance data due to their better simulation performances observed over centered log-ratio transformed data. The binary group identity is treated as the outcome variable for the lasso-based methods and the covariate variable for the DA methods. To evaluate the false positive control of different methods, we additionally utilized randomly shuffled binary group identities in the analysis.

Similar to the simulations, we applied method configurations and feature selection criteria listed in **Table S2** to perform genera selection with the original binary group labels and the randomly shuffled labels. Same random fold splits were applied to different lasso-based methods. Selected genera and total running time were collected. For each selected genus from each method using the true binary labels, we calculated the area under the ROC curve (AUC) with respect to the true binary groups.

We assess the false positive control of various methods based on the selected number of taxa using the randomly shuffled labels. Due to random shuffling, no taxa are expected to be detected as associated with the groups. Thus, any selected features can be treated as false positive findings, where the percentage of selected genera can be interpreted as the false positive rate. To visualize the results, a heatmap was produced with colors representing the false positive rates for each dataset for each method. For selected taxa based on the true binary labels, we generate heatmaps to compare the number of selected taxa, the median taxon-specific AUC, and the running time as descriptive metrics. Due to the lack of gold standard genera for each study, no inferences were made about the sensitivity of the methods.

#### 4.8.2 MSKCC allo-HCT Cohort

The 16S rRNA microbiome sequencing dataset of MSKCC patients receiving first allo-HCT between January 2009 and June 2021 was utilized to investigate the associations between genera and survival outcomes. The patient and fecal sample cohort has been partly described in past studies [29, 31, 68, 79], while the more recent samples between 2018 and 2021 were also included in analyses reported in this work. Detailed descriptions on sample collection and storage, DNA extraction, and bioinformatic pre-processing pipelines have been made available [31, 79]. Samples with sequencing depth *<* 5000 were excluded from the analysis. ASVs from the same genus were combined at genus level for subsequent analysis.

Two analysis cohorts were derived as illustrated in **Fig.S9**. We defined day 0 as the date of HCT. The peri-engraftment sample cohort consisted of the latest samples collected between day 7 and 21 relative to HCT for the 912 patients who had at least one sample collected between day 7 and 21. In contrast, the longitudinal sample cohort contained 8,967 samples from 1,415 patients, including the last sample collected prior to HCT and all samples post HCT for each patient.

Three survival endpoints of interest were defined, namely overall survival (OS), transplant-related survival (TRM), and graft-versus-host disease (GVHD)-related survival (GRM). Patients were censored at the time of last contact or at the time of second transplant, whichever occurred earlier. For TRM and GRM, we followed the hierarchical definition of competing risks [4]. Specifically, TRM or GRM will be censored by the competing risk of relapse or progression of disease. Patients who did not have recorded relapse or progression time, but with death due to relapse or disease progression, were also classified as having the endpoint of relapse or progression. For patients who did not experience relapse and progression, and also did not die due to relapse and progression, the causes of death would determine TRM and GRM. Here, TRM consists of all causes of death apart from relapse and progression, while GRM is a subset of TRM where patients died from GVHD or died after having GVHD. For the analysis associated with the peri-engraftment sample cohort, the time-to-event is landmarked at the sample collection time of the peri-engraftment sample. For the longitudinal sample cohort, the time origin is set as the time of transplant, while patients will enter the risk set at time 0 or the time of collection of the first stool sample, whichever happened earlier.

`FLORAL` was applied to investigate the association between genera and the survival endpoints defined above, adjusted for age, conditioning intensity, graft source, and disease type, using both the peri-engraftment sample cohort and the longitudinal sample cohort. The longitudinal microbial features were treated as time-dependent covariates, under the last-value-carried-forward assumption [71]. Cox proportional hazards model was applied for the OS, while Fine-Gray subdistributional proportional hazards model was applied for TRM and GRM. To assess how reliably `FLORAL` select microbial features using peri-engraftment samples versus longitudinal samples, the two-step variable selection procedure was repeated for 100 times under randomly generated 5-fold cross validation splits. For each survival endpoint, the percentages of times being selected using *λ* = *λ*_1se_ out of 100 repeated runs were compared across the peri-engraftment and the longitudinal cohorts for taxa selected at least once.

Other methods listed in **Table S2** were also applied for feature selection for OS. For lasso-based methods, the same 100 5-fold splits used for `FLORAL` were used to generate taxa selection probabilities for `glmnet` and `zeroSum`, where `glmnet` with relative abundance, log-transformed counts, and centered log-ratio transformed counts were applied for both peri-engratment and longitudinal cohorts, while `zeroSum` was only applied for the peri-engraftment cohort due to its incompatibility with time-dependent covariates. Using the OS indicators as patient group labels, the DA methods were also applied to select differentially abundant genera across the two groups with the configurations listed in **Table S2**.

## 5 Data and Code Availability

Open-source R package `FLORAL` can be accessed via GitHub (https://vdblab.github.io/FLORAL) or CRAN (https://cran.r-project.org/package=FLORAL). R scripts used for analyses can be accessed via GitHub (https://github.com/vdblab/FLORAL-analysis/). 16S rRNA sequencing datasets for the 39 studies were retrieved from https://figshare.com/articles/dataset/16S_rRNA_Microbiome_Datasets/14531724 [78]. 16S rRNA sequencing dataset for the MSKCC allo-HCT cohort can be downloaded from https://doi.org/10.6084/m9.figshare.13584986 [79].

### Algorithm 1

Iterative optimization algorithm for (8) with given *λ* and *µ*. Note that the following algorithm assumes no intercept term. The algorithm with intercept term can be derived similarly. *⊙* denotes element-wise multiplication.

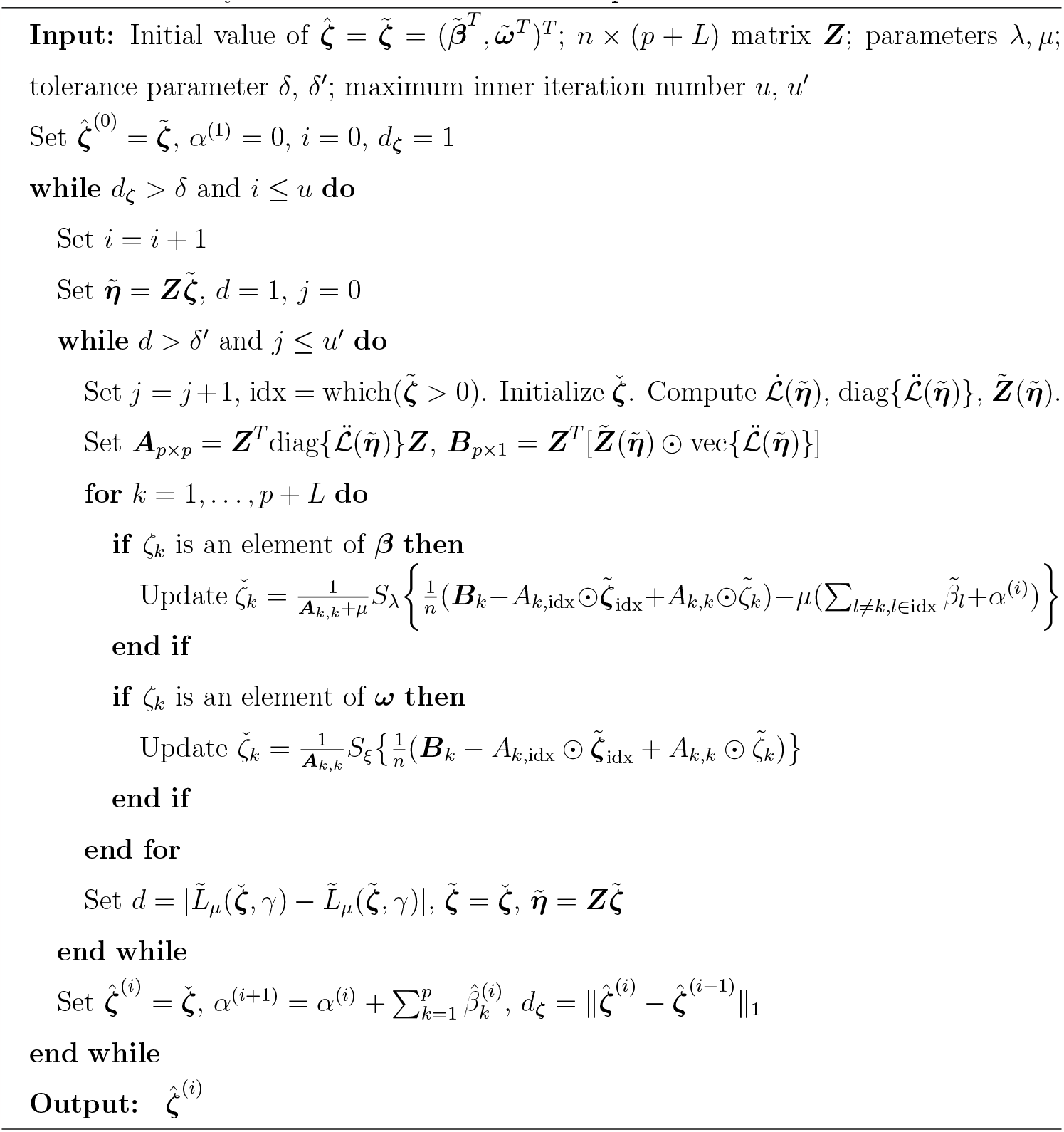

