## Supplemental Figures and Tables for "Enhanced Feature Selection for Microbiome Data using FLORAL: Scalable Log-ratio Lasso Regression"

### 1 Supplementary Figures and Tables

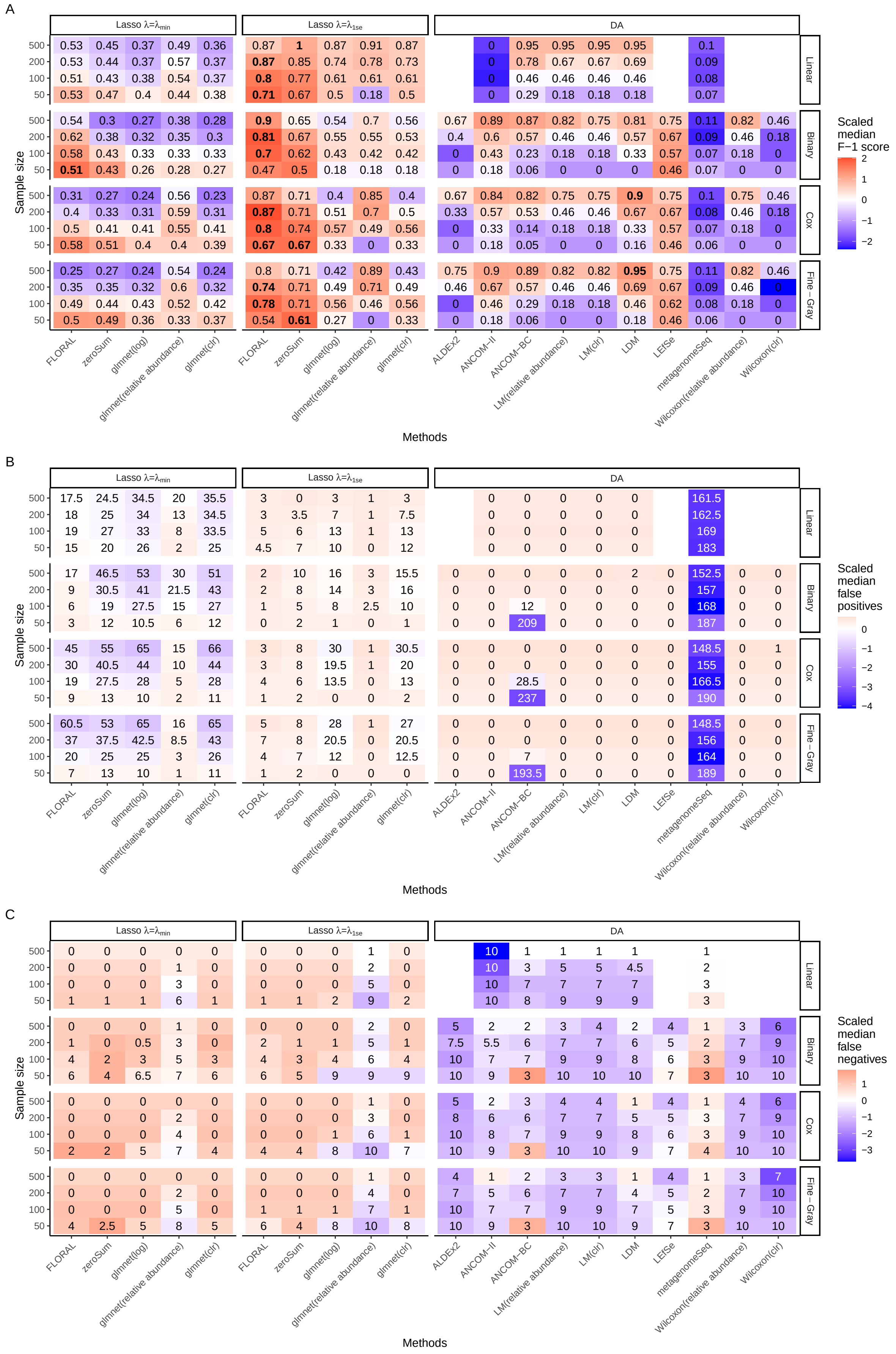
A
Lasso λ=λmin
Lasso λ=λ1se
DA
Linear
Binary
Cox
Fine−Gray
Sample size
Methods
Scaled median F−1 score
FLORAL
zeroSum
glmnet(log)
glmnet(relative abundance)
glmnet(clr)
ALDEx2
ANCOM−II
ANCOM−BC
LM(relative abundance)
LM(clr)
LDM
LEfSe
metagenomeSeq
Wilcoxon(relative abundance)
Wilcoxon(clr)
B
Scaled median false positives
C
Scaled median false negatives

Fig. S1 *(preceding page)*: **A.** median $F_1$ score, **B.** Median number of false positive features, and **C.** median number of false negative features obtained by lasso and DA methods for linear, binary, survival, and competing risks outcomes out of 100 simulations with $u = 0.5, p = 500, s = 0.8, \rho = 0$ and different choices of effect sizes $n = 50, 100, 200, 500$, where there were 10 true features in each simulation run. For each simulation scenario, scaled medians across all methods with mean zero and standard deviation one were used for color visualization. For the DA methods, the censoring indicator of the survival or competing risks outcomes were used to define patient groups except for LDM, where the Martingale residual was first computed then correlated with taxa abundances. Part of the DA methods were not evaluated for continuous outcome due to incompatibility. The adjusted p-value cutoff was set as 0.05 for all DA methods.

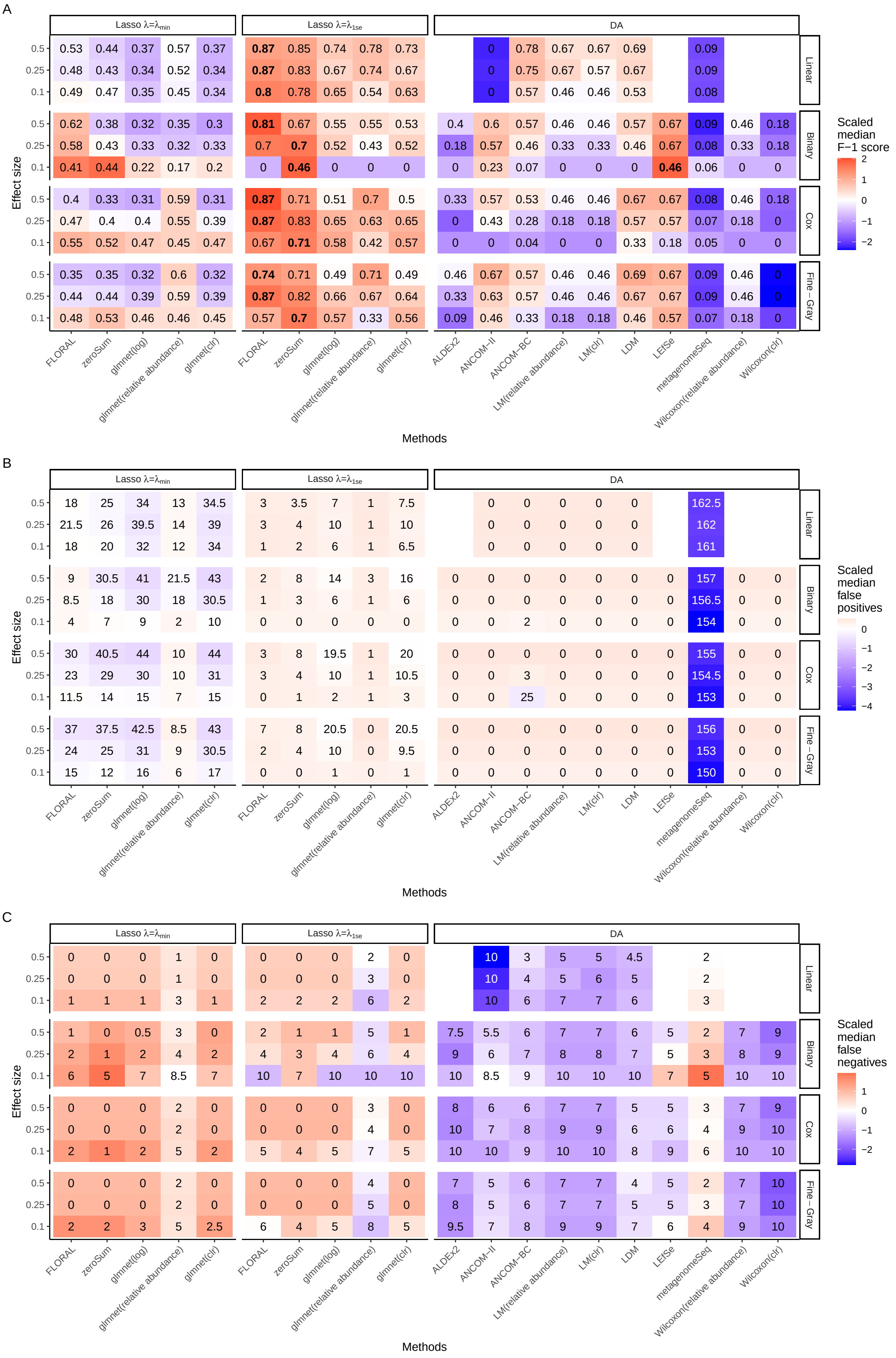
A
B
C
Lasso λ=λmin
Lasso λ=λ1se
DA
Linear
Binary
Cox
Fine−Gray
Effect size
Methods
FLORAL
zeroSum
glmnet(log)
glmnet(relative abundance)
glmnet(clr)
ALDEx2
ANCOM−II
ANCOM−BC
LM(relative abundance)
LM(clr)
LDM
LEfSe
metagenomeSeq
Wilcoxon(relative abundance)
Wilcoxon(clr)
Scaled median F−1 score
Scaled median false positives
Scaled median false negatives

Fig. S2 *(preceding page)*: **A.** median $F_1$ score, **B.** Median number of false positive features, and **C.** median number of false negative features obtained by lasso and DA methods for linear, binary, survival, and competing risks outcomes out of 100 simulations with $n = 200, p = 500, s = 0.8, \rho = 0$ and different choices of effect sizes $u = 0.1, 0.25, 0.5$, where there were 10 true features in each simulation run. For each simulation scenario, scaled medians across all methods with mean zero and standard deviation one were used for color visualization. For the DA methods, the censoring indicator of the survival or competing risks outcomes were used to define patient groups except for LDM, where the Martingale residual was first computed then correlated with taxa abundances. Part of the DA methods were not evaluated for continuous outcome due to incompatibility. The adjusted p-value cutoff was set as 0.05 for all DA methods.

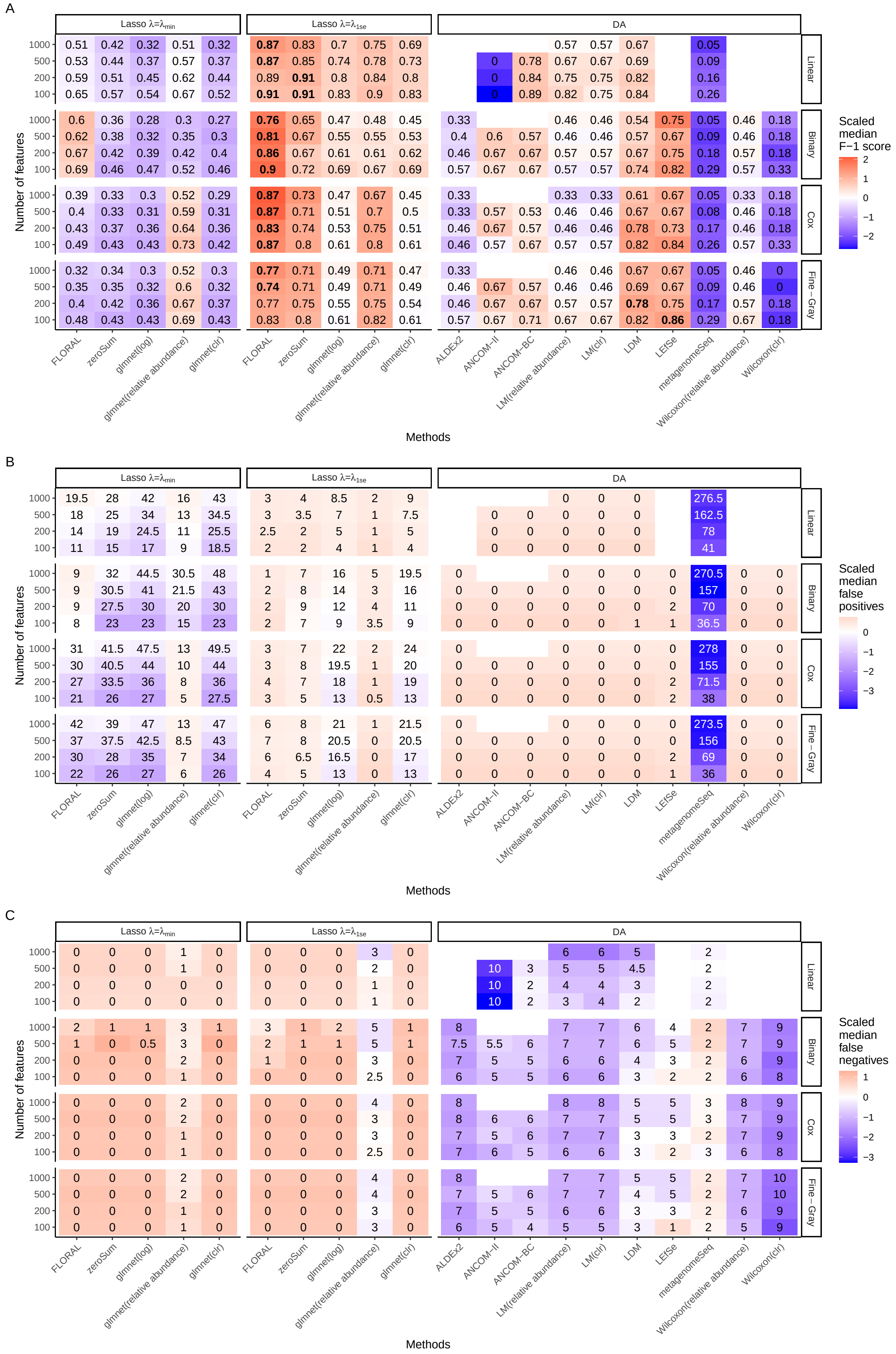
A
B
C
Lasso λ=λmin
Lasso λ=λ1se
DA
Linear
Binary
Cox
Fine−Gray
Number of features
Methods
FLORAL
zeroSum
glmnet(log)
glmnet(relative abundance)
glmnet(clr)
ALDEx2
ANCOM−II
ANCOM−BC
LM(relative abundance)
LM(clr)
LDM
LEfSe
metagenomeSeq
Wilcoxon(relative abundance)
Wilcoxon(clr)
Scaled median F−1 score
Scaled median false positives
Scaled median false negatives

Fig. S3 *(preceding page)*: **A.** median $F_1$ score, **B.** Median number of false positive features, and **C.** median number of false negative features obtained by lasso and DA methods for linear, binary, survival, and competing risks outcomes out of 100 simulations with $u = 0.5, n = 200, s = 0.8, \rho = 0$ and different choices of numbers of features $p = 100, 200, 500, 1000$. For each simulation scenario, scaled medians across all methods with mean zero and standard deviation one were used for color visualization. For the DA methods, the censoring indicator of the survival or competing risks outcomes were used to define patient groups except for LDM, where the Martingale residual was first computed then correlated with taxa abundances. Part of the DA methods were not evaluated for continuous outcome due to incompatibility. Part of the DA methods were not evaluated for $p = 1000$ due to memory overflow. The adjusted p-value cutoff was set as 0.05 for all DA methods.

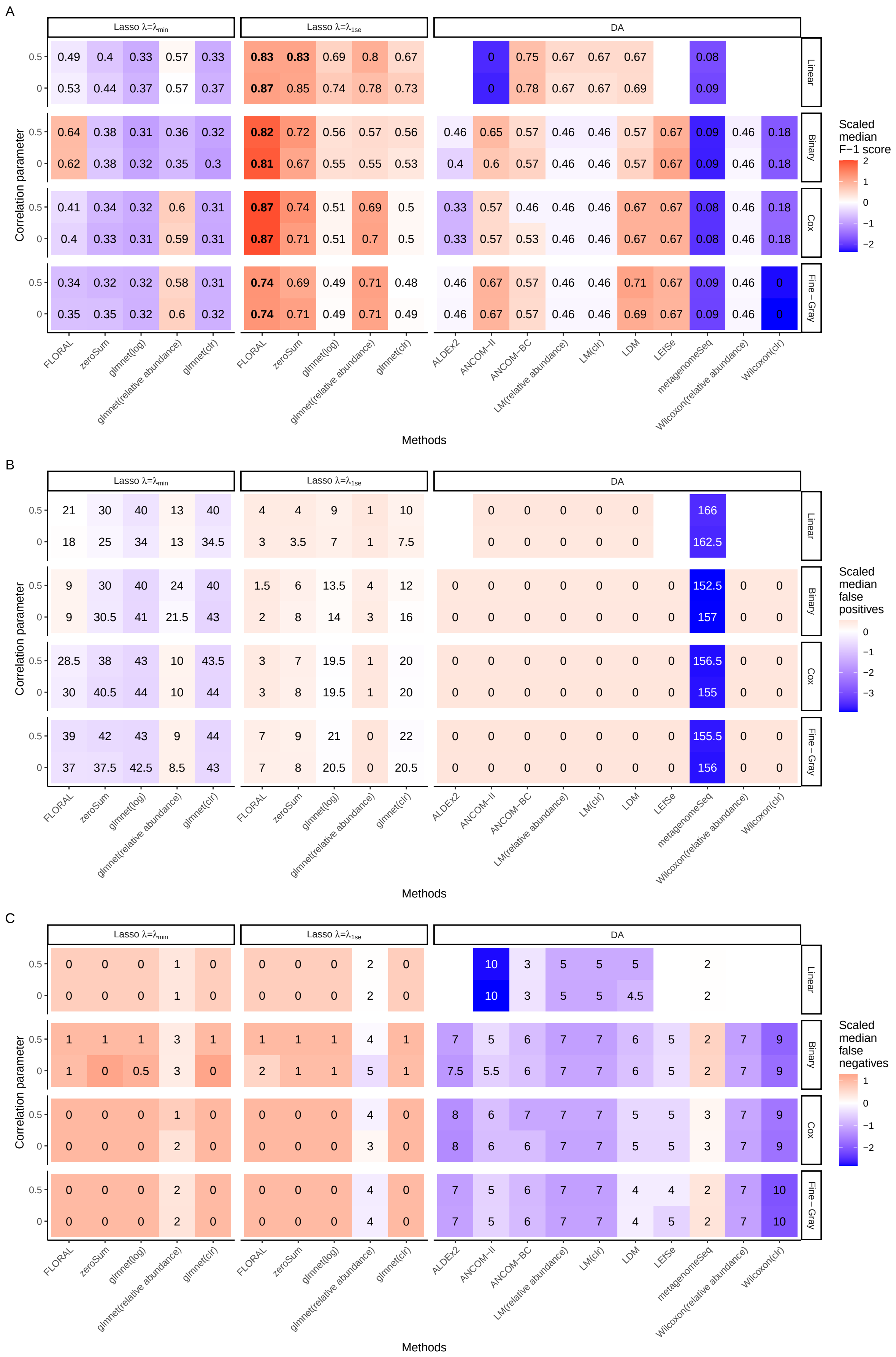
A
Lasso λ=λmin
Lasso λ=λ1se
DA
Correlation parameter
0.5
0
0.49 0.4 0.33 0.57 0.33 0.83 0.83 0.69 0.8 0.67 0 0.75 0.67 0.67 0.67 0.08
0.53 0.44 0.37 0.57 0.37 0.87 0.85 0.74 0.78 0.73 0 0.78 0.67 0.67 0.69 0.09
Linear
0.64 0.38 0.31 0.36 0.32 0.82 0.72 0.56 0.57 0.56 0.46 0.65 0.57 0.46 0.46 0.57 0.67 0.09 0.46 0.18
0.62 0.38 0.32 0.35 0.3 0.81 0.67 0.55 0.55 0.53 0.4 0.6 0.57 0.46 0.46 0.57 0.67 0.09 0.46 0.18
Binary
0.41 0.34 0.32 0.6 0.31 0.87 0.74 0.51 0.69 0.5 0.33 0.57 0.46 0.46 0.46 0.67 0.67 0.08 0.46 0.18
0.4 0.33 0.31 0.59 0.31 0.87 0.71 0.51 0.7 0.5 0.33 0.57 0.53 0.46 0.46 0.67 0.67 0.08 0.46 0.18
Cox
0.34 0.32 0.32 0.58 0.31 0.74 0.69 0.49 0.71 0.48 0.46 0.67 0.57 0.46 0.46 0.71 0.67 0.09 0.46 0
0.35 0.35 0.32 0.6 0.32 0.74 0.71 0.49 0.71 0.49 0.46 0.67 0.57 0.46 0.46 0.69 0.67 0.09 0.46 0
Fine−Gray
Scaled median F−1 score
2 1 0 −1 −2
FLORAL zeroSum glmnet(log) glmnet(relative abundance) glmnet(clr)
FLORAL zeroSum glmnet(log) glmnet(relative abundance) glmnet(clr)
ALDEx2 ANCOM−II ANCOM−BC LM(relative abundance) LM(clr) LDM LEfSe metagenomeSeq Wilcoxon(relative abundance) Wilcoxon(clr)
Methods
B
Lasso λ=λmin
Lasso λ=λ1se
DA
Correlation parameter
21 30 40 13 40 4 4 9 1 10 0 0 0 0 0 166
18 25 34 13 34.5 3 3.5 7 1 7.5 0 0 0 0 0 162.5
Linear
9 30 40 24 40 1.5 6 13.5 4 12 0 0 0 0 0 0 0 152.5 0 0
9 30.5 41 21.5 43 2 8 14 3 16 0 0 0 0 0 0 0 157 0 0
Binary
28.5 38 43 10 43.5 3 7 19.5 1 20 0 0 0 0 0 0 0 156.5 0 0
30 40.5 44 10 44 3 8 19.5 1 20 0 0 0 0 0 0 0 155 0 0
Cox
39 42 43 9 44 7 9 21 0 22 0 0 0 0 0 0 0 155.5 0 0
37 37.5 42.5 8.5 43 7 8 20.5 0 20.5 0 0 0 0 0 0 0 156 0 0
Fine−Gray
Scaled median false positives
0 −1 −2 −3
Methods
C
Lasso λ=λmin
Lasso λ=λ1se
DA
Correlation parameter
0 0 0 1 0 0 0 0 2 0 10 3 5 5 5 2
0 0 0 1 0 0 0 0 2 0 10 3 5 5 4.5 2
Linear
1 1 1 3 1 1 1 1 4 1 7 5 6 7 7 6 5 2 7 9
1 0 0.5 3 0 2 1 1 5 1 7.5 5.5 6 7 7 6 5 2 7 9
Binary
0 0 0 1 0 0 0 0 4 0 8 6 7 7 7 5 5 3 7 9
0 0 0 2 0 0 0 0 3 0 8 6 6 7 7 5 5 3 7 9
Cox
0 0 0 2 0 0 0 0 4 0 7 5 6 7 7 4 4 2 7 10
0 0 0 2 0 0 0 0 4 0 7 5 6 7 7 4 5 2 7 10
Fine−Gray
Scaled median false negatives
1 0 −1 −2
Methods

Fig. S4 *(preceding page)*: **A.** median $F_1$ score, **B.** Median number of false positive features, and **C.** median number of false negative features obtained by lasso and DA methods for linear, binary, survival, and competing risks outcomes out of 100 simulations with $u = 0.5, n = 200, p = 500, s = 0.8$ and different choices of correlation levels $\rho = 0, 0.5$, where there were 10 true features in each simulation run. For each simulation scenario, scaled medians across all methods with mean zero and standard deviation one were used for color visualization. For the DA methods, the censoring indicator of the survival or competing risks outcomes were used to define patient groups except for LDM, where the Martingale residual was first computed then correlated with taxa abundances. Part of the DA methods were not evaluated for continuous outcome due to incompatibility. The adjusted p-value cutoff was set as 0.05 for all DA methods.

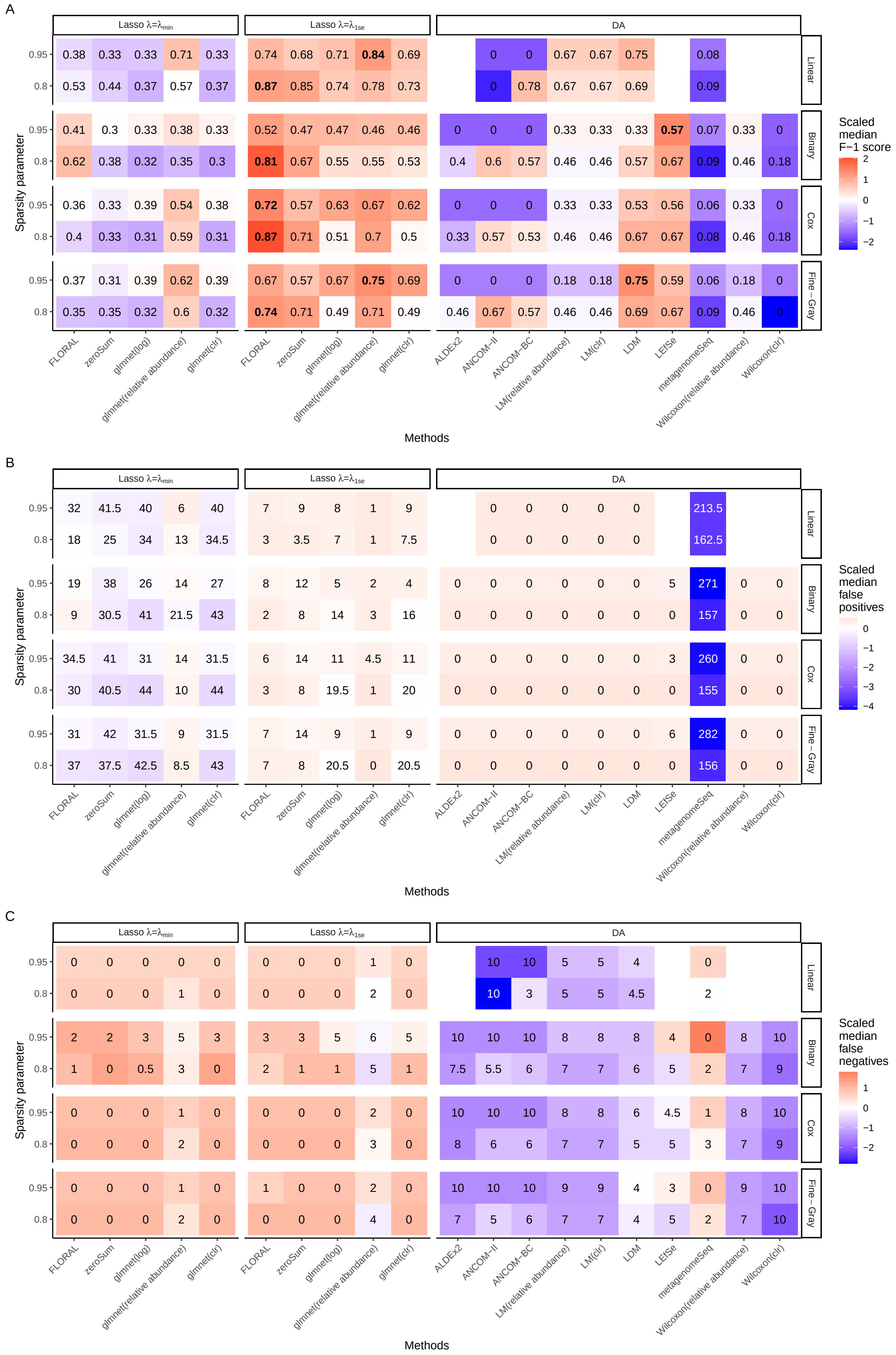
A
B
C
Lasso λ=λmin
Lasso λ=λ1se
DA
Linear
Binary
Cox
Fine−Gray
Sparsity parameter
Methods
FLORAL
zeroSum
glmnet(log)
glmnet(relative abundance)
glmnet(clr)
ALDEx2
ANCOM−II
ANCOM−BC
LM(relative abundance)
LM(clr)
LDM
LEfSe
metagenomeSeq
Wilcoxon(relative abundance)
Wilcoxon(clr)
Scaled median F−1 score
Scaled median false positives
Scaled median false negatives

Fig. S5 *(preceding page)*: **A.** median $F_1$ score, **B.** Median number of false positive features, and **C.** median number of false negative features obtained by lasso and DA methods for linear, binary, survival, and competing risks outcomes out of 100 simulations with $u = 0.5, n = 200, p = 500, \rho = 0$ and different choices of sparsity levels $s = 0.8, 0.95$, where there were 10 true features in each simulation run. For each simulation scenario, scaled medians across all methods with mean zero and standard deviation one were used for color visualization. For the DA methods, the censoring indicator of the survival or competing risks outcomes were used to define patient groups except for LDM, where the Martingale residual was first computed then correlated with taxa abundances. Part of the DA methods were not evaluated for continuous outcome due to incompatibility. The adjusted p-value cutoff was set as 0.05 for all DA methods.

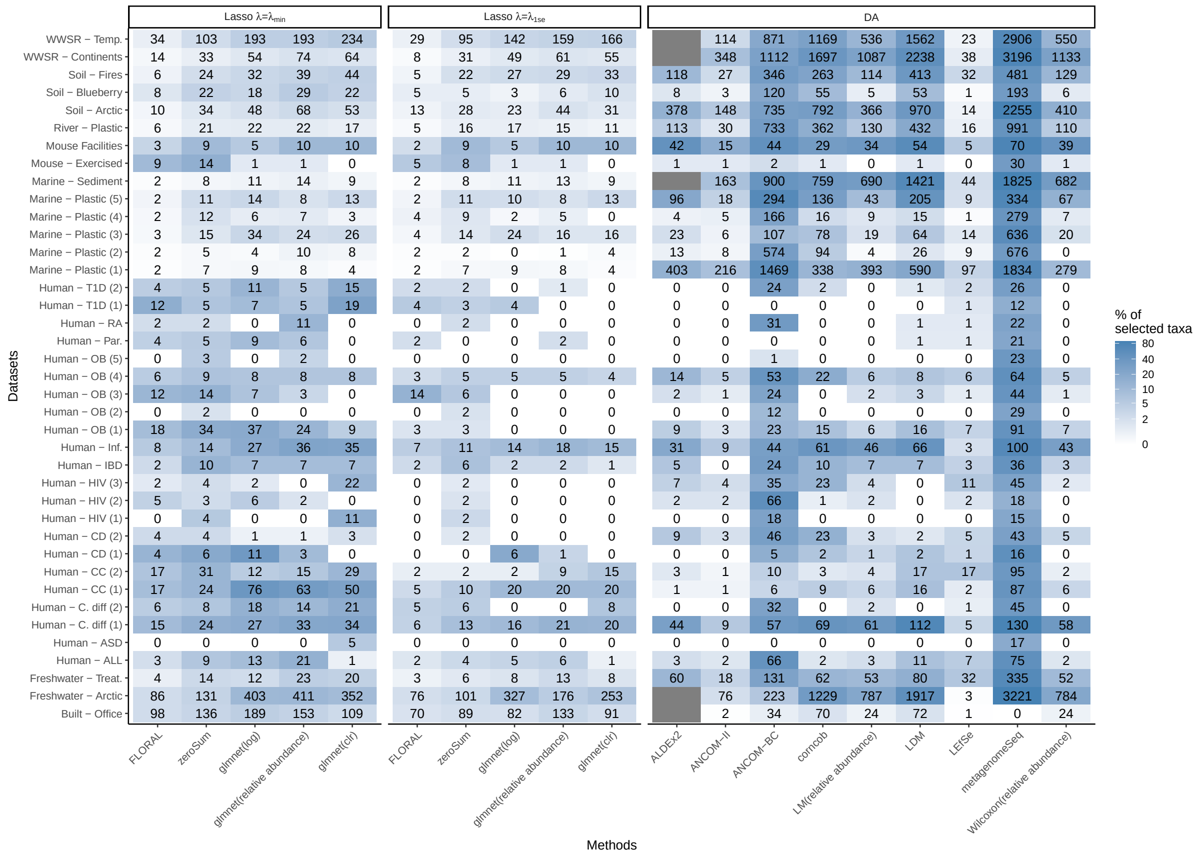


Fig. S6: Number of selected taxa from the 39 publicly available 16S microbiome data sets by feature selection methods, without group label shuffling. The color scheme represents the percentage of selected taxa out of all taxa in a certain data set. Part of data were unavailable for ALDEx2 due to memory overflow.

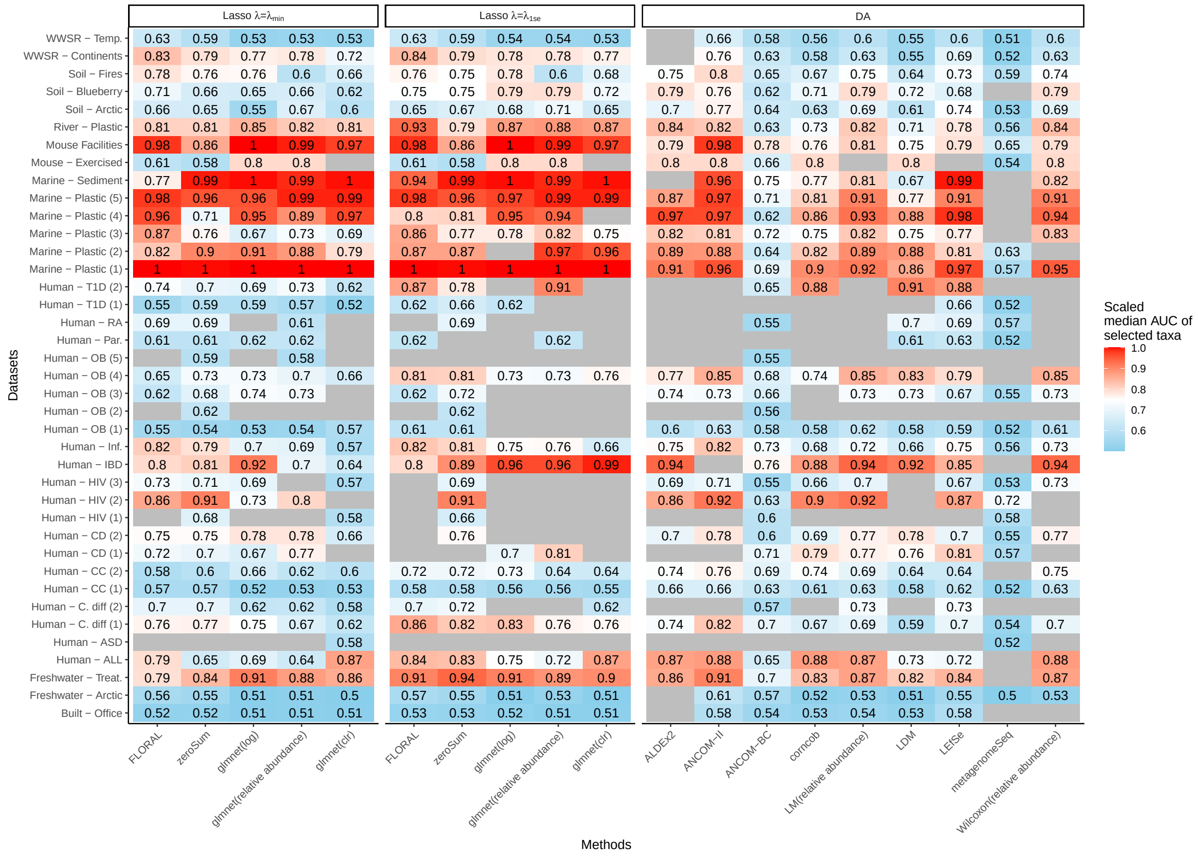


Fig. S7: Median AUC of selected features against comparison group labels from the 39 publicly available 16S microbiome data sets by feature selection methods, without group label shuffling. The median AUCs from different methods on each single data set were scaled for color visualization. Cells of grey color indicate that no features were selected. Part of data were unavailable for ALDEx2 due to memory overflow.

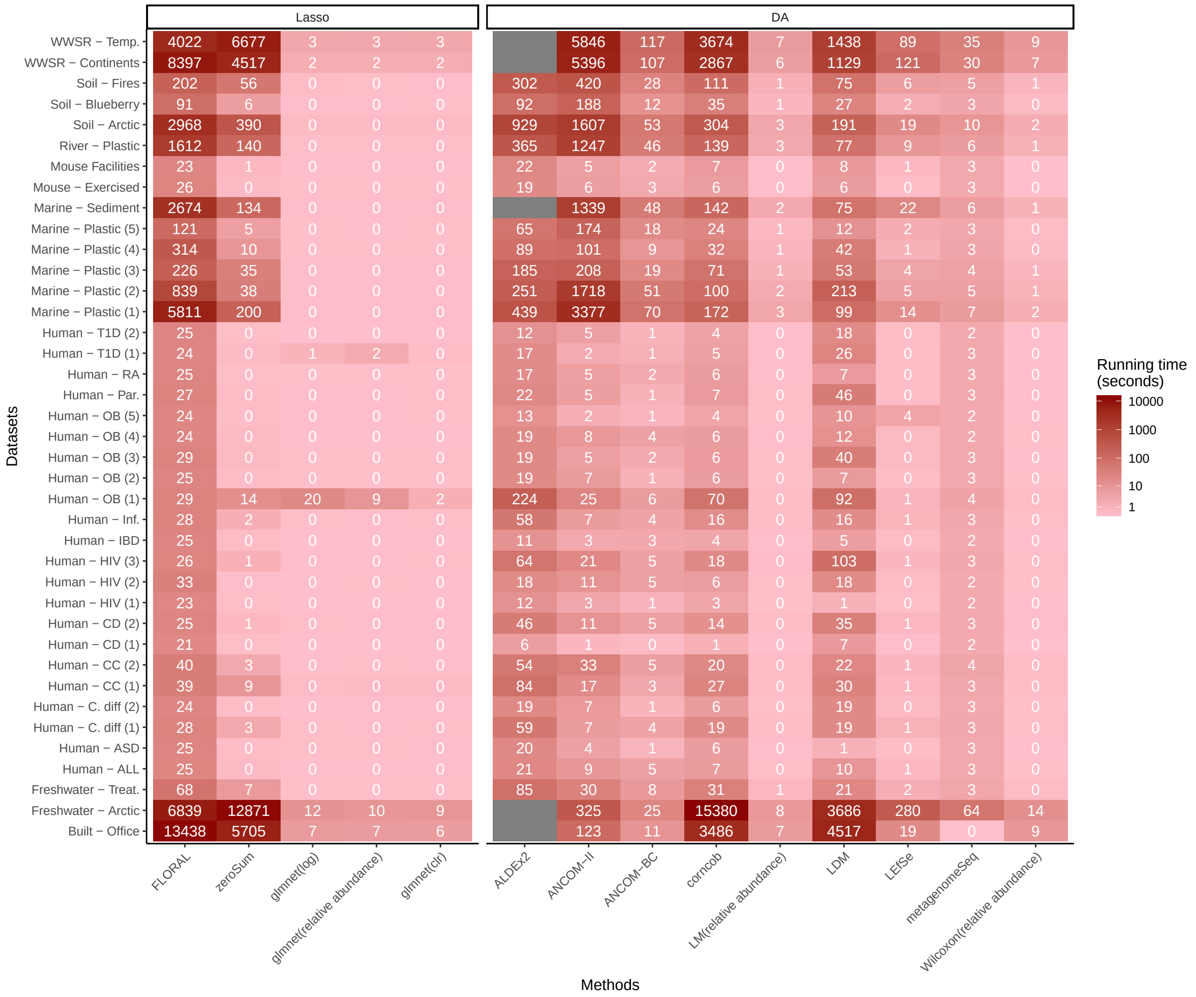


Fig. S8: Running time in seconds of different methods on the 39 publicly available 16S microbiome data sets without group label shuffling. Part of data were unavailable for ALDEx2 due to memory overflow.

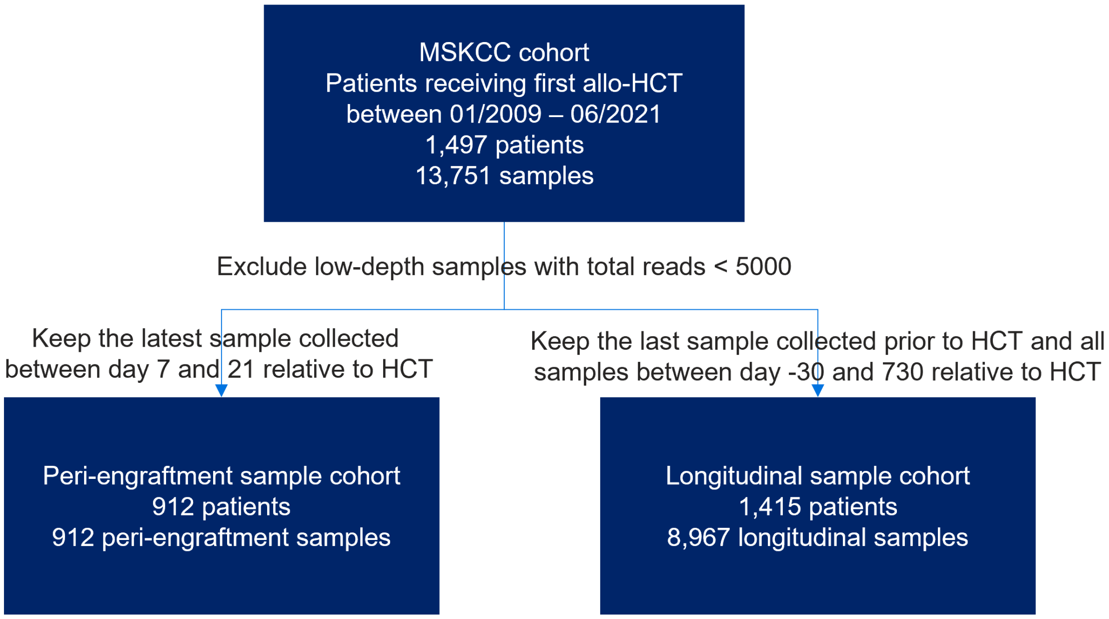


Fig. S9: Derivation of the MSKCC peri-engraftment sample cohort and longitudinal sample cohort.

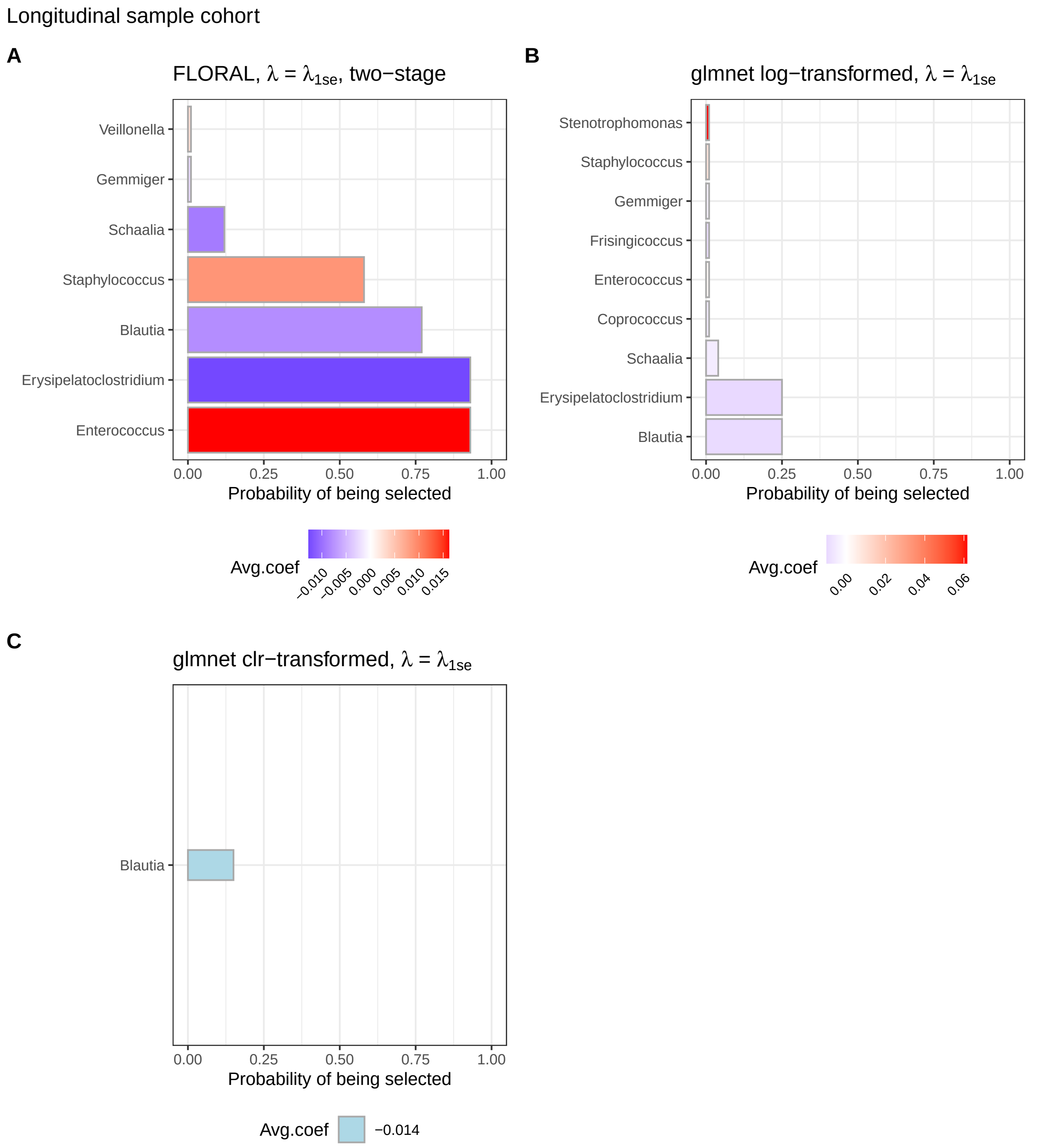


Fig. S10: Probabilities of genera being selected from 100 repeats of 5-fold cross-validation with random fold split for Cox model of overall survival with longitudinal samples using **A.** `FLORAL` with $\lambda = \lambda_{1\text{se}}$ and two-stage variable selection; **B.** `glmnet` with $\log(. + 1)$-transformed count data at $\lambda = \lambda_{1\text{se}}$; **C.** `glmnet` with centered log-ratio transformed count data at $\lambda = \lambda_{1\text{se}}$. Results from `glmnet` with relative abundances were not shown since no genera were selected in any cross validation runs at $\lambda = \lambda_{1\text{se}}$. Results from `zeroSum` were not shown due to incompatibility with time-dependent feature data. The color scheme represents the average lasso coefficient estimates of the corresponding genus at $\lambda^{(i)} = \lambda_{1\text{se}}$ over $i = 1, \ldots, 100$ repeats.

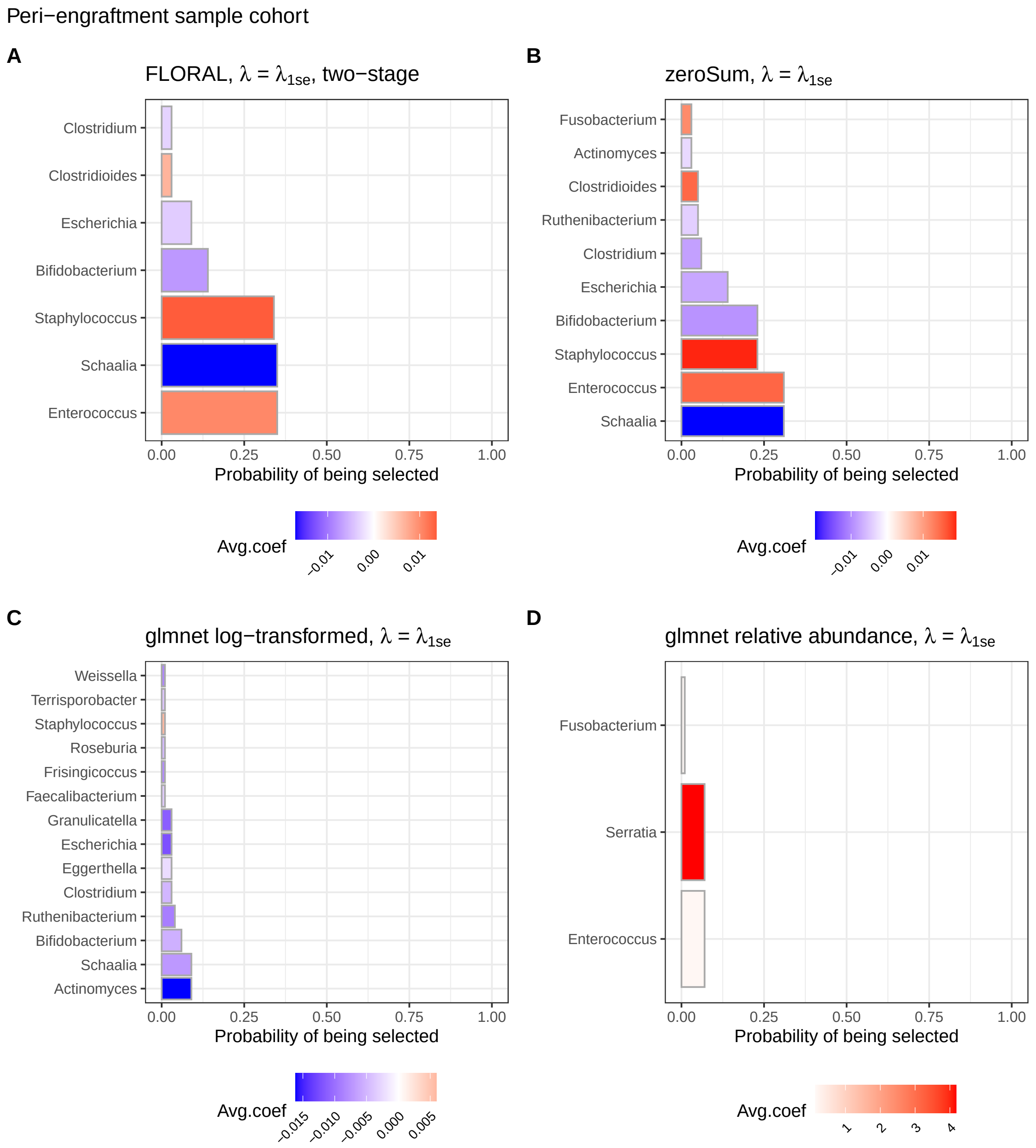


Fig. S11: Probabilities of genera being selected from 100 repeats of 5-fold cross-validation with random fold split for Cox model of overall survival with peri-engraftment samples using **A.** `FLORAL` with $\lambda = \lambda_{1\text{se}}$ and two-stage variable selection; **B.** `zeroSum` with $\lambda = \lambda_{1\text{se}}$; **C.** `glmnet` with $\log(.+1)$-transformed count data at $\lambda = \lambda_{1\text{se}}$; **D.** `glmnet` with relative abundance data at $\lambda = \lambda_{1\text{se}}$. Results from `glmnet` with centered log-ratio transformation were not shown since no genera were selected in any cross validation runs at $\lambda = \lambda_{1\text{se}}$. The color scheme represents the average lasso coefficient estimates of the corresponding genus at $\lambda^{(i)} = \lambda_{1\text{se}}$ over $i = 1, \ldots, 100$ repeats.

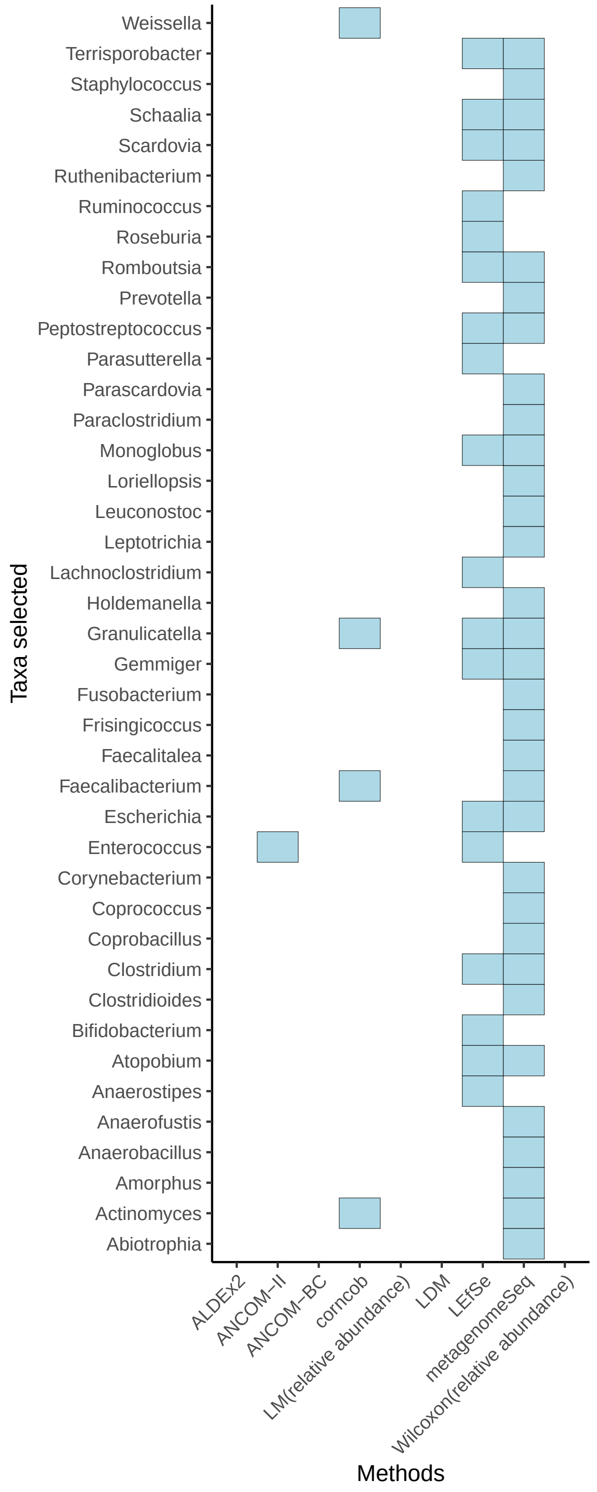


Fig. S12: Genera associated with overall survival selected by the DA methods for the MSKCC allo-HCT data. The censoring indicator of overall survival were used to define binary patient groups for all methods except for LDM, where the Martingale residual was first computed then correlated with taxa abundances. The adjusted p-value cutoff was set as 0.05 for all methods.

Table S1: Patient characteristics of the MSKCC allo-HCT cohort with peri-engraftment samples and longitudinal samples.

| **Patient Characteristic** | Patients with peri-engraftment samples **N = 912**[1] | Patients with longitudinal samples **N = 1,415**[1] |
|---|---|---|
| Disease type | | |
| AML/ALL/MDS/MPN | 679 (74%) | 1,024 (72%) |
| Others | 233 (26%) | 391 (28%) |
| Graft source | | |
| T-cell depletion | 319 (35%) | 529 (37%) |
| Unmodified | 444 (49%) | 674 (48%) |
| Cord | 149 (16%) | 212 (15%) |
| Age | 58 (47, 66) | 57 (47, 65) |
| Conditioning intensity | | |
| Ablative | 460 (50%) | 732 (52%) |
| Non-ablative | 103 (11%) | 163 (12%) |
| Reduced Intensity | 349 (38%) | 520 (37%) |
| Collection day relative to HCT | 16.0 (12.0, 19.0) | - |
| Number of longitudinal observations | - | 4 (2, 9) |
| Follow-up days landmarked at sample collection | 710 (356, 716) | - |
| Follow-up days since HCT | - | 730 (359, 730) |
| GvHD-related events | | |
| Censored | 504 (55%) | 781 (55%) |
| GvHD-related mortality | 106 (12%) | 174 (12%) |
| Relapse/progression of disease | 254 (28%) | 372 (26%) |
| Others | 48 (5.3%) | 88 (6.2%) |

[1] n (%); Median (IQR)

AML: acute myeloid leukemia; ALL: acute lymphoblastic leukemia;

MDS: myelodysplastic syndromes; MPN: myelodproliferative neoplasms;

HCT: hematopoietic cell transplantation; GvHD: graft-versus-host disease.

Table S2: Methods applied in the analyses and the corresponding configurations, feature selection criteria, and versions of R packages if applicable.

| Methods (R function) | Configurations | Selection criteria | Package version |
|---|---|---|---|
| `ALDEx2`[1] | 1000 MC samples | FDR $< 0.05$ | 1.28.1 |
| `ANCOM-II` [2] | Default | FDR $< 0.05$, W-statistic $> 0.9$ | source code |
| `ANCOM-BC` [3] | Default | FDR $< 0.05$ | 1.2.2 |
| `corncob` [4] | Default | FDR $< 0.05$ | 0.3.1 |
| `FLORAL` (This method) | Default | Proposed 2-step procedure | 0.2.0 |
| `glmnet` [5] | Default | Non-zero coefficients at $\lambda_{\text{min}}$ and $\lambda_{\text{1se}}$ | 4.1-4 |
| `LDM` [6] | Default | FDR $< 0.05$ | 4.0 |
| LEfSe [7] (`lefser`) | Default | Wilcoxon $< 0.05$, LDA $> 2$ | 1.6.0 |
| LM (`stats::lm`) [8] | Default | FDR $< 0.05$ | 4.2.1 |
| `metagenomeSeq` [9] | Default | FDR $< 0.05$ | 1.38.0 |
| Wilcoxon [8] (`stats::wilcox.test`) | Default | FDR $< 0.05$ | 4.2.1 |
| `zeroSum` [10] | Default | Non-zero coefficients at $\lambda_{\text{min}}$ and $\lambda_{\text{1se}}$ | 2.0.6 |
